# Rad51 Interaction Analysis Reveals a Functional Interplay Among Recombination Auxiliary Factors

**DOI:** 10.1101/738179

**Authors:** Bilge Argunhan, Masayoshi Sakakura, Negar Afshar, Misato Kurihara, Kentaro Ito, Takahisa Maki, Shuji Kanamaru, Yasuto Murayama, Hideo Tsubouchi, Masayuki Takahashi, Hideo Takahashi, Hiroshi Iwasaki

## Abstract

Although Rad51 is the key protein in homologous recombination (HR), a major DNA double-strand break repair pathway, several auxiliary factors interact with Rad51 to promote productive HR. Here, we present an interdisciplinary characterization of the interaction between Rad51 and Swi5-Sfr1, a widely conserved auxiliary factor. NMR and site-specific crosslinking experiments revealed two distinct sites within the intrinsically disordered N-terminus of Sfr1 that cooperatively bind to Rad51. Although disruption of this binding severely impaired Rad51 stimulation in vitro, interaction mutants did not show any defects in DNA repair. Unexpectedly, in the absence of the Rad51 paralogs Rad55-Rad57, which constitute another auxiliary factor complex, these interaction mutants were unable to promote DNA repair. Our findings provide molecular insights into Rad51 stimulation by Swi5-Sfr1 and suggest that, rather than functioning in an independent subpathway of HR as was previously proposed, Rad55-Rad57 facilitates the recruitment of Swi5-Sfr1 to Rad51.

---

DNA double-strand breaks (DSBs) are a particularly toxic form of DNA damage in which a DNA molecule is broken into two fragments. A major DSB repair pathway is homologous recombination (HR). During HR, an intact stretch of DNA that shares sequence similarity to the DSB site is identified and utilized as a template for synthesis-dependent repair. Dysregulation of HR results in misrepair of DSBs, resulting in genomic instability, a potent driver of tumorigenesis^1^.

HR is initiated by the formation of 3’ single-stranded DNA (ssDNA) at the DSB site. This ssDNA is bound by RPA then the ubiquitous RecA-family recombinase Rad51, which forms a right-handed nucleoprotein filament. The Rad51 filament is able to capture intact double-stranded DNA (dsDNA) and–by assessing the extent of base-pairing with the filamentous ssDNA–identify regions of DNA that share substantial sequence similarity to the DSB site^2^. After initial pairing with the complementary strand of the dsDNA, the Rad51 filament further displaces the noncomplementary strand by driving strand transfer, resulting in the formation of an intermediate structure known as a displacement loop. The 3’ end of the invading strand is then utilized as a primer for DNA synthesis, leading to its extension and the recovery of lost genetic information. In the simplest case, ejection of this extended strand allows it to anneal with the complementary DNA on the other side of the DSB^3^. Recombinational DNA repair is completed following gap filling by further DNA synthesis and ligation of resultant nicks.

Efforts to elucidate the underlying biochemistry of HR have typically involved measuring the ability of purified Rad51 to drive pairing and subsequent strand transfer of homologous DNA substrates, a process known as DNA strand exchange. Such experiments established RPA as a critical component of the DNA strand exchange reaction^4^. However, when RPA was added to the reaction concomitantly with Rad51, which more closely reflects the situation in vivo, this stimulatory effect was abolished. This paradoxical finding led to the discovery that other proteins known to be involved in HR serve as auxiliary factors that interact directly with Rad51 and can negate the inhibitory effect of RPA^5–11^.

Numerous distinct families of recombination auxiliary factors have been identified throughout eukaryotes, including Rad52, BRCA2, Rad54, Rad51 paralogs, Swi5-Sfr1 and the Shu complex^12^. Each group is thought to have non-overlapping roles in HR, although the mechanistic differences are yet to be elucidated^13–18^. Aside from the Shu complex, all auxiliary factors are capable of binding directly to Rad51, which is thought to be essential for their respective roles in HR^12^. Sfr1 was initially discovered in the fission yeast *Schizosaccharomyces pombe* as an interactor of Rad51, and along with Swi5, was shown to comprise an HR sub-pathway that functions independently of and in parallel to the Rad51 paralogs Rad55-Rad57^17, 19^. Subsequent biochemical reconstitutions demonstrated that substoichiometric concentrations of Swi5-Sfr1 were able to efficiently stimulate the strand exchange activity of Rad51 and Dmc1, the meiosis-specific RecA-family recombinase^20^. This enhancement of strand exchange was attributed to stabilization of the nucleoprotein filaments and stimulation of the recombinases’ ATPase activity^20–22^. Rad51-driven DNA strand exchange was recently shown to fit a three-step kinetic model with two reaction intermediates^23^. Swi5-Sfr1 enhanced transitioning of the first intermediate into the second intermediate and conversion of the second intermediate into reaction products, thus making it the only auxiliary factor known to potentiate Rad51 in both the presynaptic and synaptic phases of DNA strand exchange. These findings highlight the unique role of Swi5-Sfr1 as an HR regulator.

Limited proteolysis of Swi5-Sfr1 yielded a stable C-terminal fragment in complex with Swi5 (Swi5-Sfr1C, residues 181-299 of Sfr1)^24^, and this, along with the N-terminal half of Sfr1 (Sfr1N, residues 1-176 of Sfr1), could be stably expressed and purified^25^. Whereas Sfr1N was predicted to be intrinsically disordered^26, 27^, crystallographic analyses demonstrated that Swi5-Sfr1C forms a kinked structure^25^. Moreover, Swi5-Sfr1C was shown to stimulate Rad51-driven strand exchange by stabilizing the presynaptic filament and enhancing the ATPase activity of Rad51^25^. In contrast, Sfr1N had no direct effect on these activities but was seen to co-immunoprecipitate (co-IP) with Rad51. Such complex formation was not detected between Rad51 and Swi5-Sfr1C, despite the stimulatory effect of Swi5-Sfr1C on Rad51. Taken together with the observation that Swi5-Sfr1C was only able to stimulate Rad51 activity when present at much higher concentrations than full-length Swi5-Sfr1, these results led to a model in which Sfr1N keeps Swi5-Sfr1C anchored in close proximity to Rad51^25^.

Due to the use of truncated proteins in which entire domains were deleted^25^, it was not possible to determine whether Sfr1N has any function other than anchoring Swi5-Sfr1 to Rad51. To explore this, we employed an interdisciplinary approach to further characterize Sfr1N. We provide direct evidence that Sfr1N is intrinsically disordered and contains two sites that interact cooperatively with Rad51. Mutation of critical residues within these two sites rendered Rad51 refractory to the stimulatory effects of full-length Swi5-Sfr1, mimicking the results obtained with Swi5-Sfr1C (i.e., when the N-terminus of Sfr1 is absent), indicating that the sole function of Sfr1N is to facilitate the interaction between Swi5-Sfr1 with Rad51. Unexpectedly, and in contrast to the severely impaired Rad51 stimulation observed in vitro, these interaction mutants only showed defects in Rad51-mediated DNA repair in the absence of Rad55-Rad57, implying that these Rad51 paralogs can promote the recruitment of Swi5-Sfr1 to Rad51. Collectively, these results provide a molecular basis for Rad51 stimulation by Swi5-Sfr1 and reveal a novel interplay between recombination auxiliary factors.

## RESULTS

### Sfr1N is essential for the role of Swi5-Sfr1 in DNA repair

Since Sfr1N binds to Rad51 but does not stimulate DNA strand exchange, and Swi5-Sfr1C stimulates DNA strand exchange despite not forming a detectable complex with Rad51, it was proposed that Sfr1N functions exclusively to facilitate the interaction between Swi5-Sfr1C and Rad51^25^. However, it remained possible that Sfr1N only exerts a stimulatory effect when in the presence of Swi5-Sfr1C. To test this, strand exchange reactions containing purified Rad51 and plasmid-sized DNA substrates (Fig. 1a) were supplemented with equimolar concentrations of both Sfr1N and Swi5-Sfr1C. Even in this setting, Sfr1N did not have any stimulatory effect on DNA strand exchange (Fig. 1b,c), raising the possibility that it is dispensable for the physiological function of Swi5-Sfr1. To determine the requirement for these two modules in Rad51-dependent DNA repair, strains lacking the C-terminal or N-terminal half of Sfr1 (*sfr1N* and *sfr1C*, respectively) were constructed. Both strains showed the same sensitivity to DNA damage as a strain in which Sfr1 was completely absent (*sfr1Δ*; Fig. 1d). Furthermore, combining these truncations with *rad55Δ* sensitized cells to DNA damaging agents to the same degree as the *sfr1Δ rad55Δ* strain, which displays a complete loss of Rad51-dependent DNA repair (Supplementary Fig. 1a)^17, 19^. Sfr1N and Sfr1C were detected at comparable levels to full-length Sfr1 by immunoblotting, indicating that the sensitivity of the *sfr1N* and *sfr1C* strains is not due to a reduction in protein levels (Supplementary Fig. 1b). Furthermore, this sensitivity was not rescued by fusing Sfr1N or Sfr1C to the SV40 large T antigen nuclear localization signal, suggesting that the observed phenotype is not caused by a failure to localize to the nucleus (Supplementary Fig. 1c). Thus, although not essential for stimulation of Rad51 in vitro, Sfr1N is essential for the function of Swi5-Sfr1 in promoting Rad51-dependent DNA repair.

**Figure 1.**
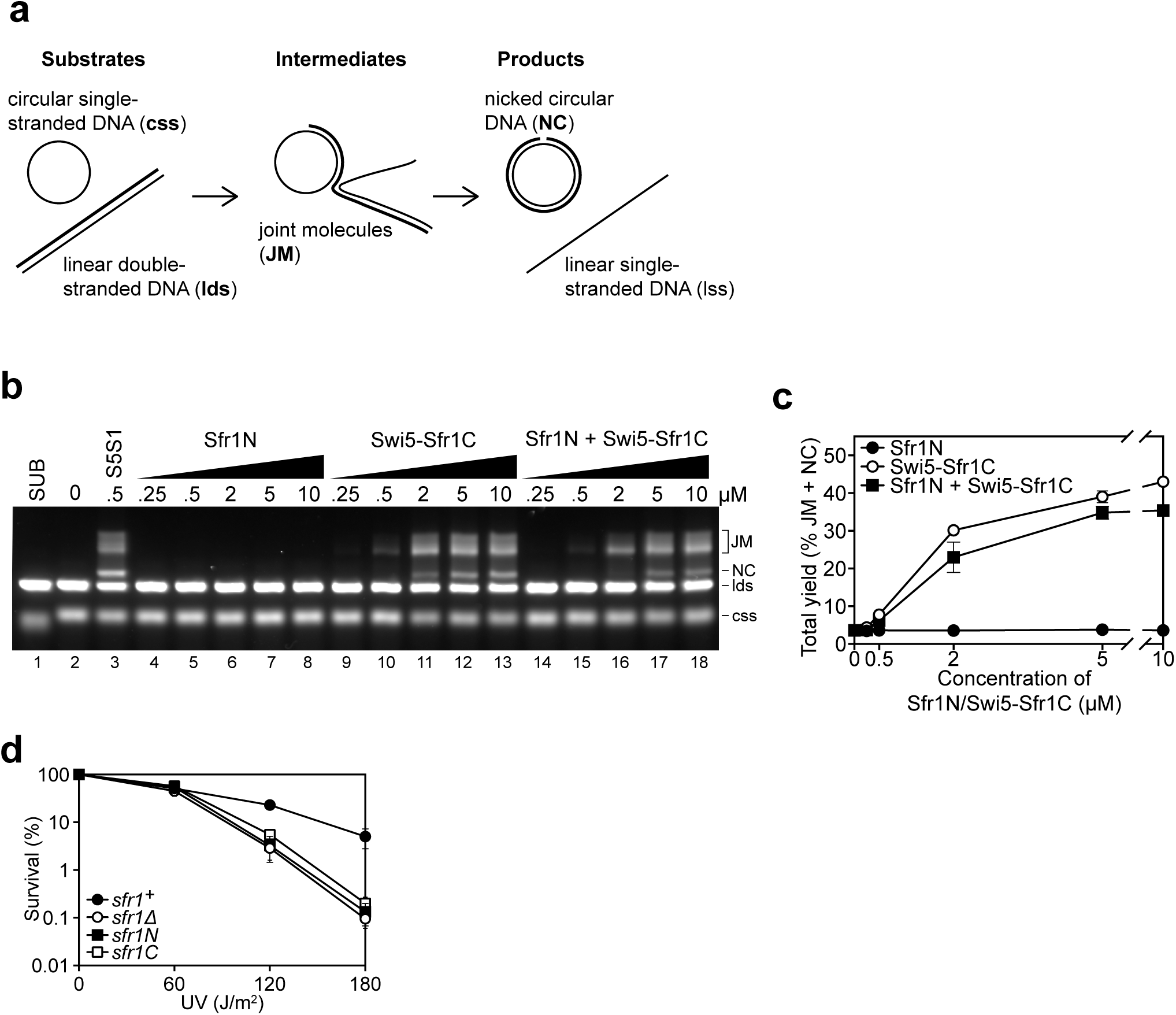
Sfr1N is essential for DNA repair mediated by Swi5-Sfr1. **(a)** Schematic of the three-strand exchange assay. **(b)** Three-strand exchange reactions containing Rad51 with the indicated concentrations of Swi5-Sfr1, or Sfr1N and/or Swi5-Sfr1C, were incubated for 2 h at 37°C and DNA was separated by agarose gel electrophoresis to visualize substrates (css, lds), intermediates (JM) and products (NC). **(c)** The percentage of DNA signal per lane corresponding to JMs and NC (total yield) was plotted against Swi5-Sfr1 concentration. **(d)** Percentage of cell survival following acute UV irradiation for the indicated strains. For **(c,d)**, mean values of three independent experiments ± s.d. are shown.

### Sfr1N comprises an intrinsically disordered and flexible domain within the Swi5-Sfr1 ensemble

Having confirmed the physiological importance of Sfr1N, a structural approach was employed to glean insights into the molecular function of Sfr1N. Primary sequence analysis and ion mobility mass spectrometry of Sfr1N suggested that this domain is intrinsically disordered^26, 27^. To directly test this, Sfr1N was analyzed by circular dichroism (CD) and nuclear magnetic resonance (NMR) spectroscopy. The CD spectrum of Sfr1N lacked local minima above 210 nm and showed a negative peak at ∼200 nm (Fig. 2a), implying a lack of secondary structural units such as α-helices and β-sheets^28^. Furthermore, examination of the ^1^H-^15^N heteronuclear single quantum coherence (HSQC) spectrum of Sfr1N revealed that most of the main chain amide protons resonated in a narrow chemical shift range between 7.7 and 8.7 ppm (Fig. 2b), which is a characteristic feature of disordered proteins^29^. To extract structural information for each residue, NMR signals from main-chain ^1^H_N_, ^13^C_α_, ^13^CO, and ^15^N_H_ atoms as well as ^13^C_β_ resonances were assigned by analyzing a set of triple resonance spectra. This was assisted by the ^1^H-^15^N HSQC spectra of selectively-^15^N labeled versions of Sfr1N and several Sfr1N variants (Supplementary Fig. 2a-c).

**Figure 2.**
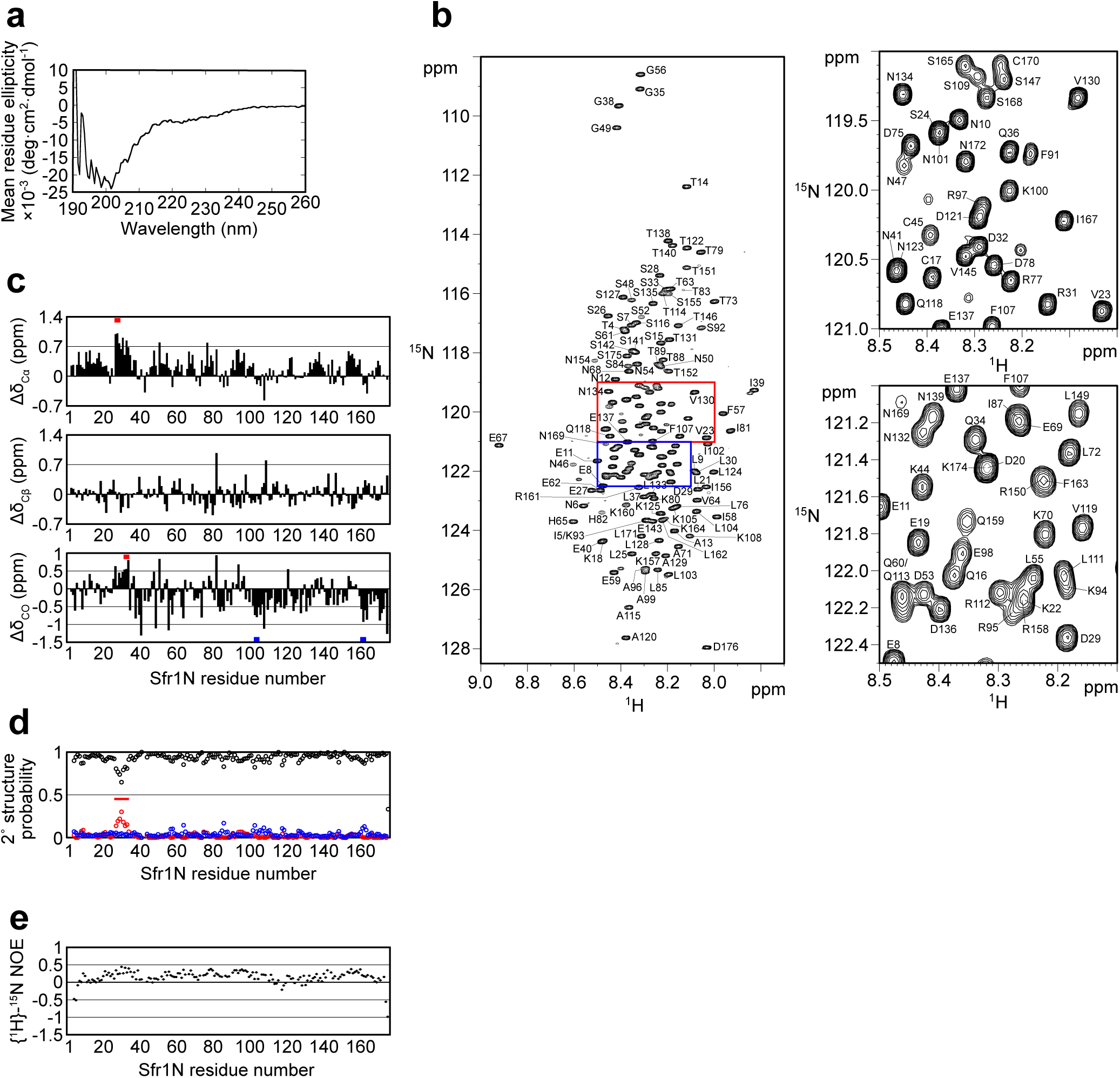
The N-terminal half of Sfr1 is an intrinsically disordered and flexible domain. **(a)** Far UV spectrum of Sfr1N. **(b)** ^1^H-^15^N HSQC spectrum of Sfr1N. Regions outlined in red and blue are enlarged in the top right and bottom right panels, respectively. **(c)** Secondary chemical shifts of ^13^C*_α_* (top), ^13^C*_β_* (middle), and ^13^CO (bottom) were obtained and plotted against the corresponding position in Sfr1N. Values that suggest potential *α*-helix and *β*-strand formation are indicated with red and blue bars, respectively. **(d)** Talos+ prediction of secondary structure probabilities. Black, red, and blue circles indicate the probabilities of random coils, *α*-helices, and *β*-strands, respectively. **(e)** {^1^H}-^15^N heteronuclear NOE was measured for Sfr1N. All analyzed main chain NHs showed NOE values of less than 0.44.

The chemical shifts obtained for the main-chain and ^13^C_β_ atoms enabled secondary structure prediction. The secondary chemical shift of ^13^C_α_, ^13^C_β_, and ^13^CO atoms, which is the chemical shift difference between the measured values and the corresponding amino acids in random coil peptides, was determined (Fig. 2c). The majority of Sfr1N residues showed ^13^C secondary chemical shift values within a limited range, suggesting that random coil structures are present in these regions. Nonetheless, a few groups of residues exhibited secondary chemical shift values outside of this range, raising the possibility that some structures resembling α-helices (E27 to D29, D32 to Q34) or β-strands (L103 to K105, R161 to K164) may form within Sfr1N^30^. Further secondary structure analysis was performed using the program TALOS+^31^, which predicted Sfr1N to be entirely disordered, with a low probability for α-helix formation from E27 to S33 (Fig. 2d).

To analyze the dynamical features of Sfr1N, the steady state heteronuclear nuclear Overhauser effect (NOE) for the main-chain amide groups of the protein was analyzed^32, 33^. NOE values for all residues was less than 0.44, indicating that the entire protein is flexible with pico-to-nanosecond timescale motions (Fig. 2e). Such fast motions are typically observed for unstructured proteins/domains, in agreement with the above results. The NOE values were not completely uniform, with residues E27 to S33 showing slightly increased values, consistent with the possibility that this region of the protein may form an α-helix. Collectively, these results demonstrate that, unlike the structured Swi5-Sfr1C complex^25^, the N-terminal half of Sfr1 is intrinsically disordered and flexible.

### Two sites within Sfr1N interact with Rad51

Previous results indicated that Sfr1N facilitates the interaction between Swi5-Sfr1 and Rad51^25^. To identify the site(s) within Sfr1N that binds to Rad51, NMR spectra of ^15^N-labeled Sfr1N were analyzed in the absence and presence of Rad51. Superimposed ^1^H-^15^N HSQC spectra of ^15^N-labeled Sfr1N with increasing amounts of Rad51 were constructed (Fig. 3a). The most prominent spectral changes, defined as a reduction in signal intensity of >80%, were observed for 19 out of 142 non-overlapped residues (Fig. 3b,e). In addition to these marked changes, 18 and 20 residues experienced signal intensity reductions of 60-80% and 40-60%, respectively (Fig. 3e). The signal intensity of these residues was further attenuated by the incremental addition of Rad51 (Supplementary Fig. 3a-c). Most of these attenuated signals did not display obvious chemical shift changes following Rad51 binding. However, five residues (A71, T73, D75, L76, and T146) displayed incremental chemical shift changes and reductions in signal intensity upon addition of increasing amounts of Rad51 (Fig. 3c and Supplementary Fig. 3d). The remaining 60 residues that were analyzed experienced minimal effects upon addition of Rad51 (<40% reduction in signal intensity; Fig. 3d,e). These findings implicate two sites in Sfr1N, Site 1 (S84 to T114) and Site 2 (T152 to S165), where the most significantly attenuated signal intensities are sandwiched by moderately attenuated signal intensities (Fig. 3e), as being important for the Sfr1N-Rad51 interaction. Site 1 is highly basic and hydrophobic compared to other regions of Sfr1N. Positively charged residues are also a prominent feature of Site 2, but this site is not especially hydrophobic. These results suggest that, while both Sites 1 and 2 are involved in electrostatic interactions with Rad51, Site 1 may also participate in hydrophobic interactions with Rad51.

**Figure 3.**
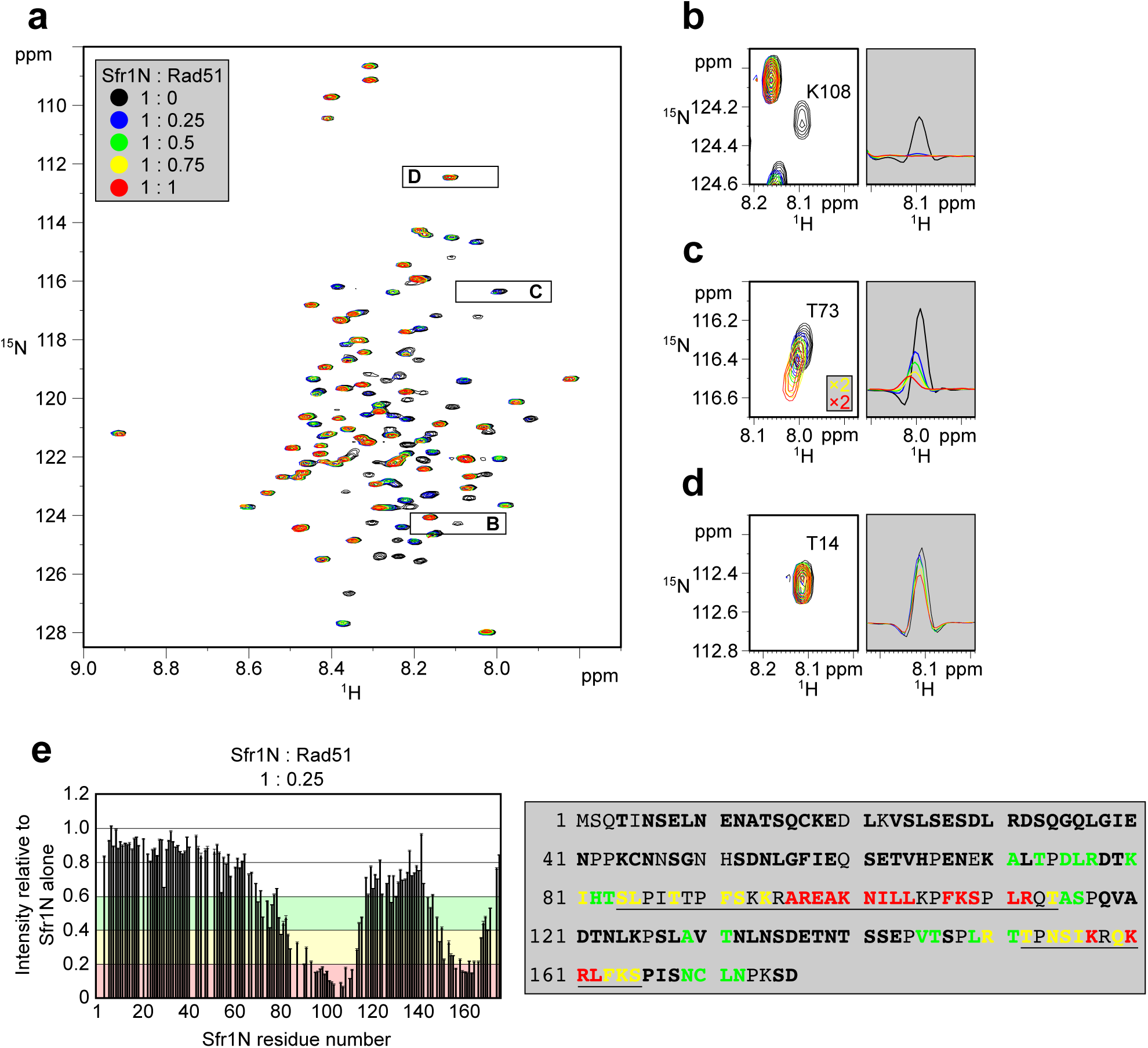
Two sites within Sfr1N interact with Rad51. **(a)** Superimposed ^1^H-^15^N HSQC spectra of ^15^N-labeled Sfr1N in the absence of Rad51 or in the presence of Rad51 at the indicated molar ratios. **(b-d)** Enlarged signals from K108, T73, and T14 (left), with slices along the ^1^H axis (right). **(e)** Signal intensity ratio of Sfr1N residues in the presence of Rad51 to those in the absence of Rad51 (left) and features of the Sfr1N amino acid sequence (right). Reductions in signal intensity of 40-60% (green), 60-80% (yellow), and >80% (red) are highlighted, along with the corresponding residues. Underlined residues correspond to Site 1 (S84 to T114) and Site 2 (T152 to S168), where the most significant signal attenuations were observed. Residues not in bold include prolines, unassigned residues, and residues with overlapped signals.

To provide further support that Sites 1 and 2 within Sfr1N interact with Rad51, site-specific crosslinking experiments were conducted (Fig. 4a). Several residues within Sites 1 and 2 were replaced with the photoreactive amino acid *p*-benzoyl-L-phenylalanine (*p*BPA) by utilizing *Escherichia coli* with an expanded genetic code^34^. Following exposure to UV light, proteins within ∼3 Å of *p*BPA become crosslinked to it^35^. Such crosslinked proteins can be detected as slow-migrating species by immunoblotting and are implicated in forming part of the interface in a protein-protein interaction^36^. Rad51 was co-expressed in *E. coli* with Sfr1N and cells were irradiated with UV. Proteins were then analyzed by immunoblotting with anti-Rad51 and anti-Sfr1 antibodies. Some non-specific crosslinking was observed when cells were treated with UV (Supplementary Fig. 3e). In addition to these non-specific crosslinks, numerous instances of specific crosslinking–defined as being dependent on a TAG mutation, the inclusion of *p*BPA in the media, and UV treatment–were observed (Fig. 4b). Whereas several positions within Site 1 showed robust crosslinking to Rad51, positions within Site 2 showed little crosslinking (Fig. 4c and Supplementary Fig. 3f), suggesting that the associations between Site 2 and Rad51 are more transient than those involving Site 1. Taken together, the results obtained from the NMR interaction analysis and site-specific crosslinking experiments indicate that two sites within Sfr1N, designated as Sites 1 and 2, interact with Rad51.

**Figure 4.**
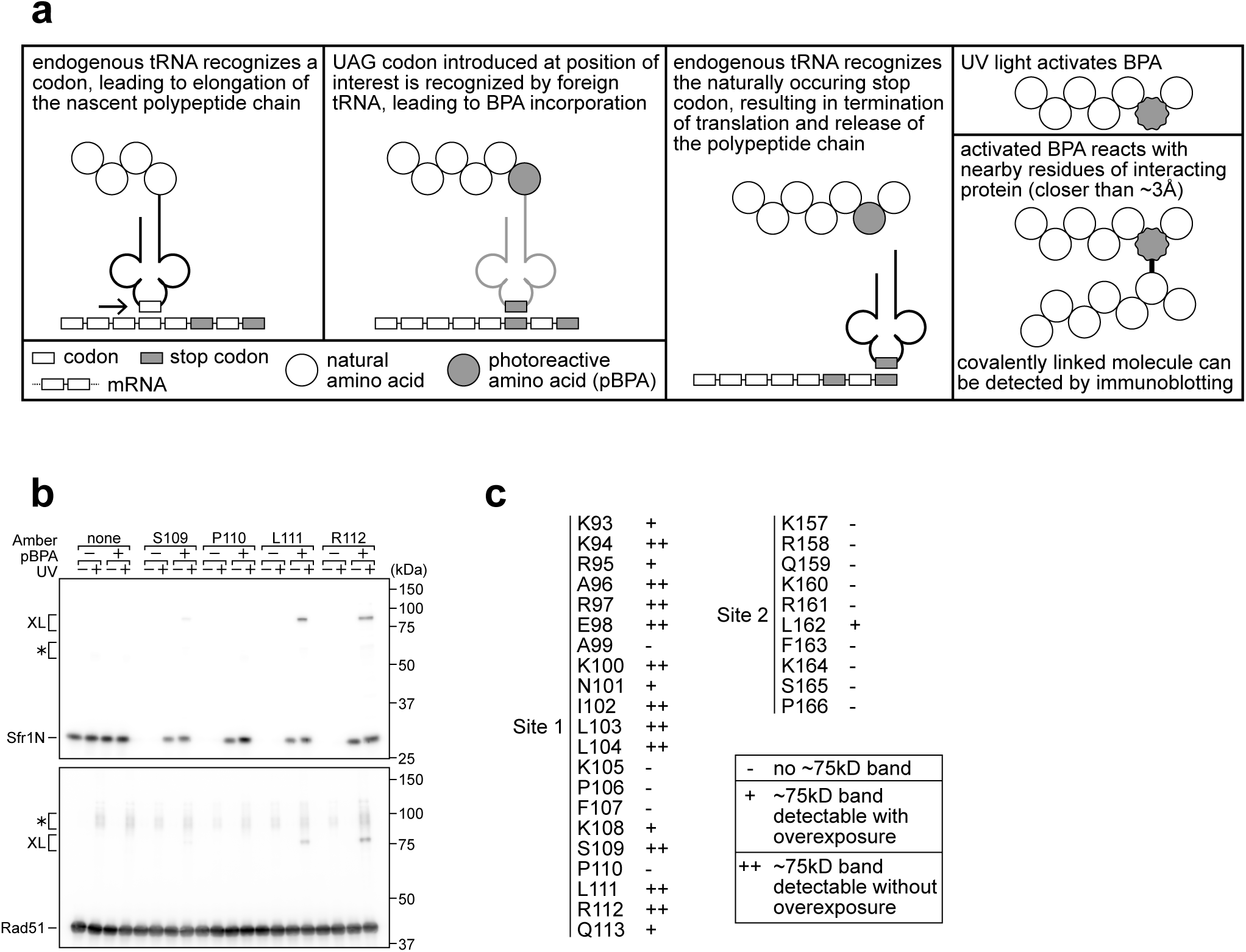
Residues within Sites 1 and 2 can be site-specifically crosslinked to Rad51. **(a)** Schematic of the crosslinking assay. **(b)** Example of Sfr1N-Rad51 crosslinking. The indicated residues were replaced with an amber codon, or the amber codon was omitted as a negative control (“none”). XL, specific crosslinks. *, non-specific crosslinks. **(c)** Summary of crosslinking results for all residues examined. See Supplementary Fig. 3 for all blots.

### Sites 1 and 2 cooperatively facilitate the physical and functional interaction between Swi5-Sfr1 and Rad51

A sequence alignment of Sfr1 orthologs within the genus *Schizosaccharomyces* highlighted conserved patches of positively charged residues in Sites 1 and 2 (Supplementary Fig. 4a). Combined with the knowledge that Sfr1N only co-IPs with Rad51 under low-salt conditions^25^, it seemed plausible that these residues might be important for electrostatic interactions with Rad51. Hence, three residues in Site 1 were mutated (3A) and four residues in Site 2 were mutated (4A). Additionally, these mutants were combined to generate the 7A mutant (Fig. 5A).

**Figure 5.**
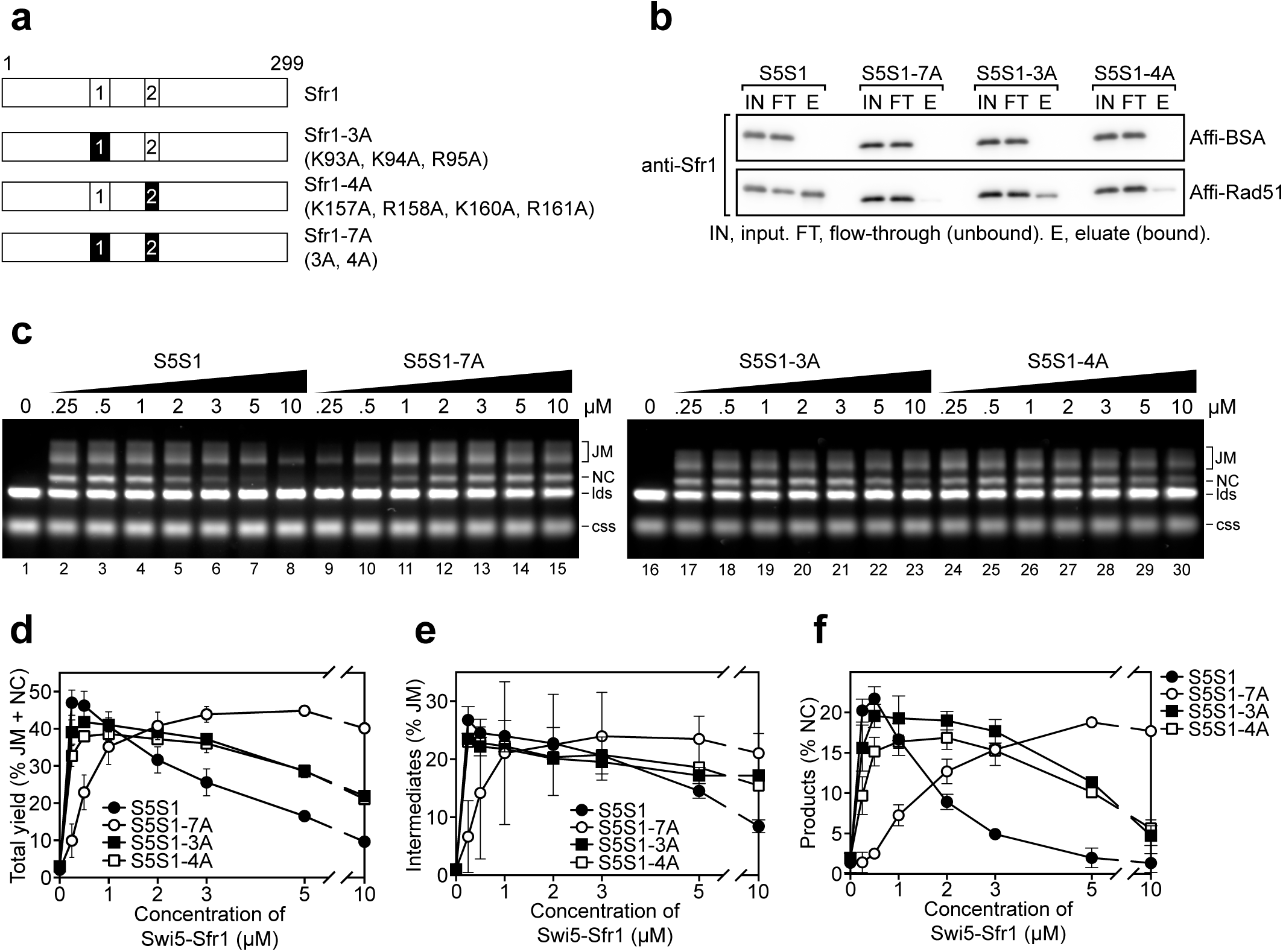
Interaction of Sites 1 and 2 with Rad51 is important for stimulation of Rad51-driven strand exchange. **(a)** Schematic of Sfr1-Rad51 interaction mutants. **(b)** The interaction of Swi5-Sfr1 (S5S1, wild type or mutants) with Rad51 was investigated by a pull-down assay. Affi-BSA is a control for nonspecific binding. **(c)** Three-strand exchange reactions were performed with the indicated concentrations of Swi5-Sfr1 (wild type or mutants). The percentage of DNA signal per lane corresponding to total yield **(d)**, JM **(e)** or NC **(f)** was plotted against Swi5-Sfr1 concentration. For **(d-f)**, mean values of three independent experiments ± s.d. are shown.

To directly assess whether these mutations disrupt the interaction with Rad51, Swi5 and full-length Sfr1 were co-purified to homogeneity (Supplementary Fig. 4b). Next, purified Rad51 was crosslinked to Affi-gel matrix and mixed with Swi5-Sfr1. A substantial fraction of wild type Swi5-Sfr1 was recovered in the eluate (Rad51-bound fraction), although some of the protein remained in the flow-through (unbound fraction; Fig. 5b). In contrast, the amount of 3A and 4A mutant proteins detected in the eluate was reduced, with much of the protein remaining in the flow-through. The 7A mutant protein was barely detected in the eluate, indicating that the binding seen in the 3A mutant was dependent on Site 2 and the binding seen in the 4A mutant was dependent on Site 1. Comparable trends were observed in a co-IP assay that did not involve crosslinking of Rad51 (Supplementary Fig. 5c). Taken together, these results suggest that Sites 1 and 2 facilitate the binding of Swi5-Sfr1 to Rad51 in a cooperative manner.

In the 7A mutant, both Sites 1 and 2 are mutated but the remainder of the N-terminus is intact. Thus, the 7A mutant can be employed to test whether the N-terminal domain of Sfr1 has any significant role other than to facilitate binding to Rad51. We therefore proceeded to characterize the biochemical activities of the 7A mutant. The 3A and 4A mutants were included to glean further insights into the nature of Rad51 stimulation by Swi5-Sfr1.

While Swi5-Sfr1C can stimulate Rad51 activity despite the absence of Sfr1N, 5-to-10-fold more of the complex is required to achieve the same level of stimulation as full-length Swi5-Sfr1 (Fig. 1b lanes 3 and 12)^25^, suggesting that the interaction between Sfr1N and Rad51 is important for efficient stimulation of strand exchange. Consistent with the observed interaction defect, substoichiometric concentrations of the 7A mutant failed to efficiently stimulate Rad51-driven strand exchange, with a substantially higher concentration of mutant protein required to achieve a wild type level of joint molecules (JMs, reaction intermediates) and nicked-circles (NCs, reaction products; Fig. 5c). At 0.25 µM, the defect of the 7A protein was more pronounced for NCs (∼15-fold reduction) than JMs (∼5-fold reduction), suggesting that the ability of Swi5-Sfr1 to stimulate both the initial pairing of homologous DNA and the subsequent strand transfer by Rad51 are defective when the interaction with Sites 1 and 2 are ablated (Fig. 5d-f). In contrast, the 3A and 4A mutants were able to promote efficient JM and NC formation at substoichiometric concentrations. Nevertheless, the loss of Rad51 stimulation observed at higher concentrations of wild type Swi5-Sfr1 was attenuated in the 3A and 4A mutants (Fig. 5c lanes 6, 21 and 28), suggesting that this loss of stimulation occurs due to unproductive interactions between Swi5-Sfr1 and Rad51. Consistent with this notion, efficient stimulation of Rad51 was maintained at higher concentrations of the 7A mutant (Fig. 5c lanes 8 and 15) and Swi5-Sfr1C (Fig. 4b,c)^25^. Collectively, these results indicate that interactions between Rad51 and both Sites 1 and 2 are important for efficient stimulation of strand exchange.

### Rad51 filament stabilization and ATPase stimulation is mediated by Sites 1 and 2

To determine why stimulation of Rad51-driven strand exchange is inefficient when Sites 1 and 2 are mutated, the molecular roles of Swi5-Sfr1 were considered. At substoichiometric concentrations, Swi5-Sfr1 effectively stabilizes Rad51 filaments^21^. Thus, it seemed feasible that the observed impairment in strand exchange might be explained by defects in Rad51 filament stabilization. To test this possibility, filament stability was examined by fluorescence anisotropy. When Rad51 binds to a fluorescently-labelled oligonucleotide and forms a filament, the fluorescence anisotropy increases due to a retardation in the motion of the labelled oligonucleotide (Fig. 6a). The dissociation of Rad51 is accompanied by a reduction in anisotropy, with the rate of decline reflective of Rad51 filament stability. Rad51-ssDNA filaments were formed in the presence of ATP and filament collapse was induced via dilution into reaction buffer containing ATP but lacking DNA and protein. In the absence of Swi5-Sfr1, the decrement in anisotropy was sharp and reached a value that was observed in the absence of protein (∼0.1) within ∼500 seconds. Inclusion of wild type Swi5-Sfr1 resulted in a slower reduction in anisotropy, indicating that the Rad51 filament had been stabilized (Fig. 6b). Strikingly, inclusion of the 7A mutant did not result in any obvious filament stabilization (Fig. 6c). Furthermore, although both 3A and 4A mutants showed some stabilization of Rad51 filaments, the magnitude of this stabilization was less than that observed for wild type protein (Fig. 6d,e). Consistent with these observations, the reaction rate constants (*k_off_*) for dissociation of Rad51-ssDNA complexes showed a substantial decline in the presence of Swi5-Sfr1, a lesser decline for the 3A and 4A mutants, and only a marginal decline for the 7A mutant (Fig. 6f). Taken together, these results indicate that Sites 1 and 2 within Sfr1N interact cooperatively with Rad51 to facilitate filament stabilization by Swi5-Sfr1.

**Figure 6.**
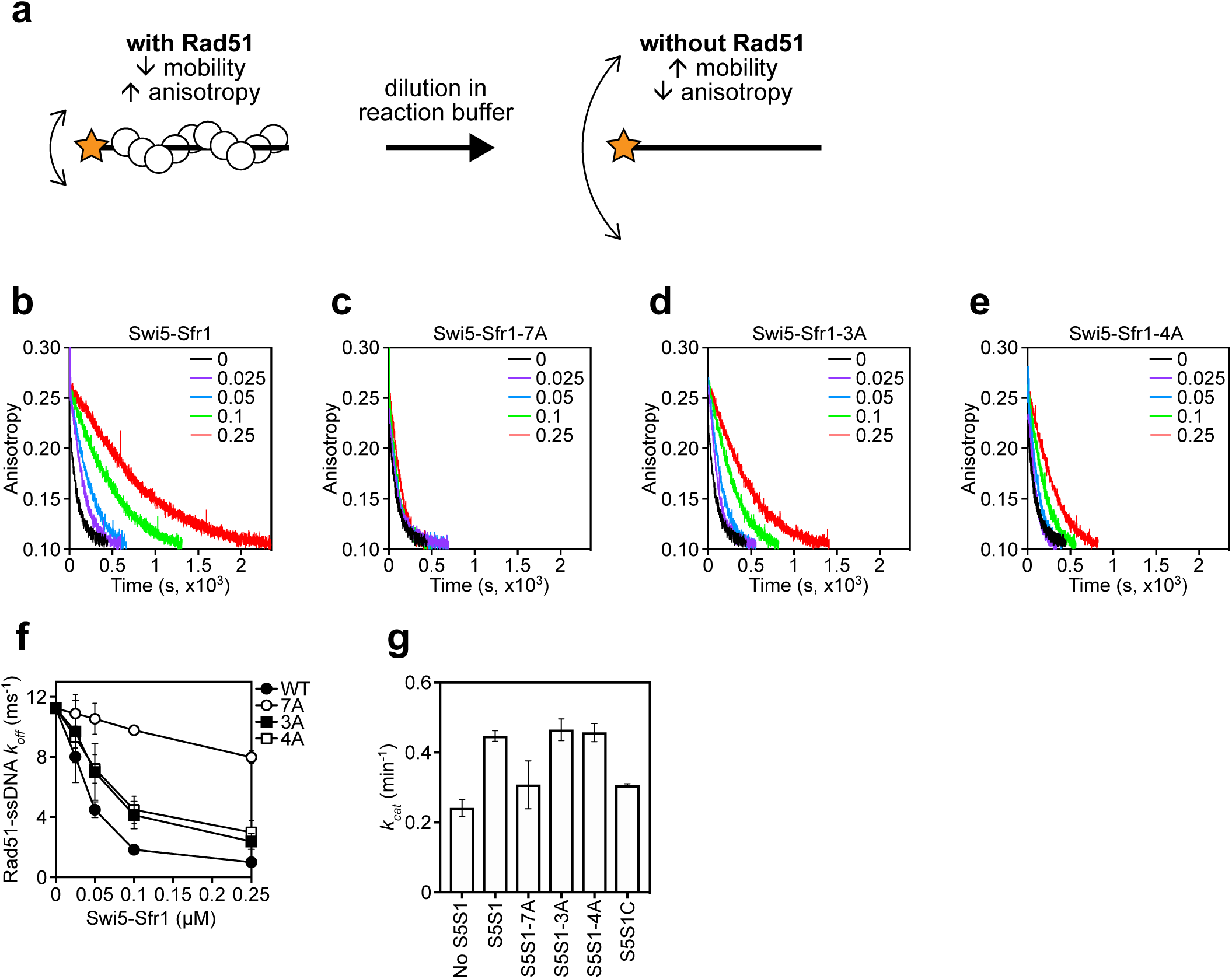
Rad51 filament stabilization and ATPase stimulation requires interactions with Sites 1 and 2. **(a)** Schematic of the fluorescence anisotropy assay. Rad51 monomers (white circles) are depicted forming a filament on an oligonucleotide (black line) labeled with the TAMRA fluorophore (orange star). **(b-e)** The anisotropy of fluorescently labelled ssDNA in complex with Rad51 was monitored following induction of filament collapse in the presence of Swi5-Sfr1 (wild type or mutants) at the indicated concentrations (µM). **(f)** *k_off_* values were determined for Rad51-ssDNA complexes at the indicated concentrations of Swi5-Sfr1. **(g)** Rad51-dependent ATP turnover was determined in the presence of the indicated Swi5-Sfr1 variant. For **(f,g)**, mean values of three independent experiments ± s.d. are shown.

In addition to stabilizing Rad51 filaments, Swi5-Sfr1 has been shown to stimulate the ATPase activity of Rad51, which is also important for efficient strand exchange^20, 21, 23^. Since substoichiometric concentrations of Swi5-Sfr1C failed to efficiently stimulate the ATPase activity of Rad51^25^, we sought to determine whether Rad51-dependent ATP hydrolysis was potentiated by the 7A mutant. As expected, wild type Swi5-Sfr1 was able to efficiently enhance ATP hydrolysis by Rad51 at substoichiometric concentrations (Swi5-Sfr1:Rad51 ratio of 1:20), with a 1.85-fold increase in ATP turnover (Fig. 6g). In contrast, the 7A mutant only managed to enhance the ATPase activity of Rad51 1.28-fold, which is similar to the 1.27-fold stimulation observed with Swi5-Sfr1C. The 3A and 4A mutants stimulated ATP hydrolysis like wild type. These results suggest that interaction of either Site 1 or 2 with Rad51 is sufficient to promote efficient stimulation of ATP hydrolysis.

### DNA repair in Sfr1-Rad51 interaction mutants is facilitated by Rad51 paralogs

Collectively, the in vitro defects of the 7A mutant are highly reminiscent of observations made with Swi5-Sfr1C (i.e., in the absence of Sfr1N). Since the *sfr1C* strain was as sensitive to DNA damage as *sfr1Δ* (Fig. 1d and Supplementary Fig. 1a,c), it seemed likely that strains expressing Sfr1 mutant proteins defective in the interaction with Rad51 would also be sensitive to DNA damage. To test this prediction, strains were constructed in which the native *sfr1^+^* gene was replaced with either *sfr1-7A*, *sfr1-3A* or *sfr1-4A*. Unexpectedly, the interaction mutants did not show any obvious sensitivity to DNA damage (Fig. 7a and Supplementary Fig. 5a). A marginal sensitivity was observed for the *sfr1-7A* strain but this was not statistically significant (Fig. 7b).

**Figure 7.**
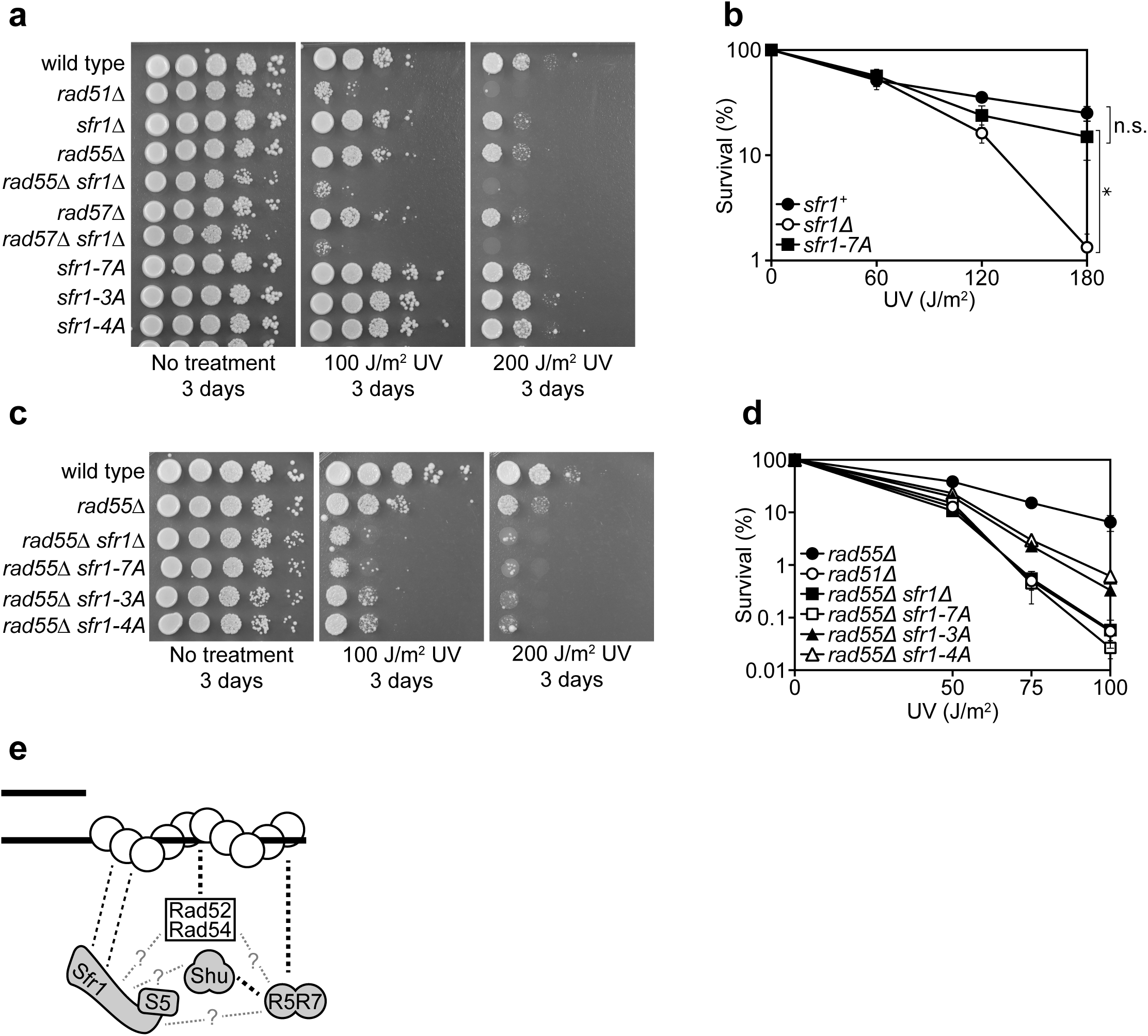
Rad55-Rad57 facilitates Swi5-Sfr1-dependent DNA repair. **(a,c)** 10-fold serial dilutions of the indicated strains were spotted onto rich media with or without acute UV treatment and grown at 30°C. **(b,d)** Percentage of cell survival was determined after acute exposure to UV. **(e)** Molecular bridging model. Rad51 monomers (white circles) are shown as a filament on ssDNA (solid black lines). Dashed black lines represent physical interactions with line thickness portraying interaction strength. Dashed grey lines represent potential physical interactions. Swi5-Sfr1 is recruited to Rad51 via two redundant mechanisms, one requiring its direct interaction with Rad51 and the other requiring Rad55-Rad57. See Discussion for details. For **(b,d)**, mean values of three independent experiments ± s.d. are shown. For **(b)**, statistical analysis was by one-way ANOVA followed by Tukey’s multiple comparisons test. *, *P* < 0.05. n.s., not significant (*P* > 0.05).

To explain these results, we considered the possibility that, although the interaction between Rad51 and Swi5-Sfr1-7A is compromised in vitro, Swi5-Sfr1-7A may still associate with Rad51 in vivo via another Rad51-interacting protein. Previous genetic studies suggested that there are two HR subpathways in *S. pombe*: one pathway is dependent on Swi5-Sfr1 and the other pathway is dependent on the Rad51 paralogs Rad55-Rad57^17, 19^. Although these two pathways are thought to function in parallel, promoting Rad51 activity independently of each other, it remained formally possible that Rad55-Rad57, which interacts with Rad51^11^, could bind to Swi5-Sfr1-7A and serve as a molecular bridge to facilitate Rad51 stimulation. To examine this possibility, the interaction mutants were introduced into the *rad55Δ* background. Strikingly, in the absence of Rad55, the *sfr1-7A* mutant showed the same DNA damage sensitivity as the *sfr1Δ* mutant (Fig. 7c). Furthermore, both the *sfr1-3A* and *sfr1-4A* mutants were more sensitive to DNA damage than *sfr1^+^* in the *rad55Δ* background, although this sensitivity was not as severe as that observed for the *sfr1Δ rad55Δ* and *sfr1-7A rad55Δ* strains (Fig. 7c,d). Similar results were obtained in the *rad57Δ* background (Supplementary Fig. 5b), verifying the requirement for an intact Rad55-Rad57 auxiliary factor complex. Importantly, these results suggest that defects in the direct interaction between Swi5-Sfr1 and Rad51 are suppressed by a Rad55-Rad57-dependent mechanism that allows Swi5-Sfr1 to stimulate Rad51 in vivo.

## DISCUSSION

In this study, we characterized the interaction between Rad51, the key protein in HR, and Swi5-Sfr1, a widely conserved recombination auxiliary factor. The N-terminal half of Sfr1 was found to be essential for the role of Swi5-Sfr1 in promoting Rad51-dependent DNA repair (Fig. 1). This domain was shown to be intrinsically disordered (Fig. 2) and contain two sites that interact with Rad51 (Fig. 3,4). Although mutation of the two interaction sites disrupted the physical and functional interaction with Rad51 in vitro (Fig. 5,6), unexpectedly, defects in DNA repair were only observed in the absence of Rad55-Rad57, another auxiliary factor complex (Fig. 7).

### Cooperative interactions with Rad51 are the primary function of Sfr1N

Although Sfr1N is not essential for stimulation of Rad51-driven DNA strand exchange (Fig. 1b,c)^25^, it was found to be essential for the promotion of Rad51-dependent DNA repair by Swi5-Sfr1 (Fig. 1d and Supplementary Fig. 1a-c). NMR interaction analysis revealed that two domains within the N-terminus of Sfr1 interact with Rad51 (Fig. 3 and Supplementary Fig. 3). Mutation of Site 1 or 2 weakened the interaction with Rad51 while mutation of both sites resulted in a near-complete loss of interaction (Fig. 5b and Supplementary Fig. 4c), indicating that Sites 1 and 2 of Sfr1N bind cooperatively to Rad51. Interestingly, Rad51-driven strand exchange and ATP hydrolysis were significantly impaired only when both sites were mutated (7A mutant, Fig. 5c-f and 6g). These results indicate that the reduced interaction in the single site mutants (3A or 4A) is sufficient for Swi5-Sfr1 to fully stimulate Rad51 in these assays. Although this points towards some functional redundancy, it is possible that these assays were not sensitive enough to detect marginal defects in the stimulation of Rad51-driven strand exchange and ATP hydrolysis. Indeed, the fluorescence anisotropy assay revealed a severe defect in Rad51 filament stabilization for the 7A mutant and a modest defect for the 3A and 4A mutants (Fig. 6b-f), indicating that interaction of both Sites 1 and 2 with Rad51 is important for efficient filament stabilization. Swi5-Sfr1 has also been shown to stabilize Rad51 filaments against the F-box helicase Fbh1^37^. It would be interesting to test whether the interaction mutants can function in a similar capacity.

The reduction in NMR signals from Sites 1 and 2 in the presence of Rad51 (Fig. 3e), combined with the gradual chemical shifts observed for some residues (Fig. 3c,d), indicated that the association and dissociation of Sfr1N and Rad51 is fast on the NMR timescale, suggesting that the Sfr1N-Rad51 interaction is relatively weak. Like all RecA-family recombinases, *S. pombe* Rad51 is predicted to exist as a multimer in solution. Consistently, size-exclusion chromatography yielded a broad elution profile with a retention time corresponding to ∼160 kDa (Supplementary Fig. 6a). While the size of monomeric Rad51 (∼40kD) is too small to cause severe line-broadening of NMR signals, the interaction of multimeric Rad51 with Sfr1N could explain the drastic reduction in NMR signals for Sites 1 and 2. The NMR interaction analysis was largely substantiated by site-specific crosslinking of residues within Sites 1 and 2 to Rad51 (Fig. 4 and Supplementary Fig. 3e,f). Since both sites are involved in electrostatic interactions with Rad51, the more robust crosslinking of Site 1 may be due to the added contribution of hydrophobic interactions between Site 1 and Rad51. Alternatively, it is possible that the incorporation of the aromatic *p*BPA into Site 1 was better tolerated due to the more hydrophobic nature of this site.

Since in vitro results obtained with the 7A mutant complex closely resemble what was observed in the complete absence of Sfr1N (i.e., with Swi5-Sfr1C), we conclude that the function of Sfr1N in promoting Rad51-dependent DNA repair primarily involves cooperative binding of Sites 1 and 2 to Rad51. We nevertheless note that the 7A complex displayed a loss of DNA binding and 4A showed impaired DNA binding (Supplementary Fig. 6b). *sfr1-3A* is more sensitive to DNA damage than *sfr1^+^* in the *rad55Δ* background (Fig. 7c,d), despite DNA binding by 3A being comparable to wild type, arguing that DNA binding by Swi5-Sfr1 is not relevant to DNA repair. Furthermore, despite their difference in DNA binding, 3A and 4A were indistinguishable in all other aspects (Fig. 5-7). Although we cannot completely rule out the possibility that the defects of the 7A mutant are related to a defect in DNA binding, these results strongly suggest that DNA binding by Swi5-Sfr1 is impertinent to its role in stimulating Rad51 activity or promoting Rad51-dependent DNA repair. Mouse Swi5-Sfr1 (mSwi5-Sfr1), which stimulates Rad51 through similar mechanisms (see below), does not display any DNA binding^38^ while human Sfr1 (hSfr1) has been implicated in transcriptional regulation^39^, raising the possibility that the DNA binding activity of *S. pombe* Swi5-Sfr1 may have some relevance to a function other than DNA repair.

### Rad51 paralogs promote Swi5-Sfr1-dependent DNA repair

While the *sfr1Δ* and *rad55Δ* single mutants are sensitive to DNA damage, neither is as sensitive as *rad51Δ*, which is epistatic to both^9, 19^. However, because the *sfr1Δ rad55Δ* double mutant shows the same sensitivity as *rad51Δ* (e.g., Fig. 7a)^17, 19^, it was concluded that two independent sub-pathways of HR exist in *S. pombe*^19^. Despite the numerous defects observed in vitro, the *sfr1-7A* mutant strain was proficient for DNA repair, but this repair was completely dependent on Rad55-Rad57 (Fig. 7a-d), indicating that a Rad55-Rad57-dependent mechanism overcomes defects in the binding of Swi5-Sfr1 to Rad51.

To explain these results, we propose that the interaction of Swi5-Sfr1 with Rad51 is enabled by two redundant mechanisms: one through a direct interaction and the other through Rad55-Rad57, which interacts with Rad51^11^ and acts as a molecular bridge to facilitate the recruitment of Swi5-Sfr1 to Rad51 (Fig. 7e). Hence, although Swi5-Sfr1-7A cannot interact directly with Rad51, Rad55-Rad57 aids the recruitment of Swi5-Sfr1-7A to Rad51, allowing it to exert a stimulatory effect on Rad51; this explains why *sfr1-7A* is proficient for DNA repair. However, in the absence of Rad55 or Rad57, this tethering is lost but Swi5-Sfr1 can nevertheless promote some DNA repair via its direct interaction with Rad51, thus explaining why *rad55Δ* is not as sensitive as *rad51Δ*. It is only when both interaction mechanisms are defective, as in the *rad55Δ sfr1-7A* strain, that the promotion of Rad51-mediated DNA repair by Swi5-Sfr1 is completely lost. We therefore surmise that, while the Swi5-Sfr1 and Rad55-Rad57 sub-pathways are capable of operating independently of each other, as observed in the *rad55Δ* and *sfr1Δ* backgrounds, Swi5-Sfr1 and Rad55-Rad57 likely collaborate to promote Rad51-dependent DNA repair in wild type cells. While physical evidence for this model is lacking, Rad55-Rad57 facilitates recruitment of the Shu complex to Rad51 by binding to both and acting as a molecular bridge^14, 40^, so it could plausibly fulfill a similar role for Swi5-Sfr1. However, unlike the Shu complex, Swi5-Sfr1 can interact directly with Rad51 (Fig. 5b and Supplementary Fig. 4c)^20, 25^, so any contribution made by Rad55-Rad57 to this interaction would enhance rather than enable complex formation with Rad51. The requirement for such a mechanism may stem from the fact that the direct interaction between Swi5-Sfr1 and Rad51 is relatively weak (see above). Indeed, previous attempts by us and others to co-IP Swi5-Sfr1 and Rad51 from yeast extracts has been unsuccessful^19, 41^, suggesting that the cellular interaction is too weak and/or transient to capture. It is tempting to speculate that Rad55-Rad57, Swi5-Sfr1, and the Shu complex exist as a higher-order auxiliary factor complex, perhaps as part of a Rad52-containing DNA repair center^42, 43^. Evidence for the existence of such a complex and elucidation of its molecular function will be the focus of future research.

### Evolutionary conservation of Sfr1N structure and function

CD and NMR spectroscopy directly demonstrated that Sfr1N is an intrinsically disordered and dynamic domain within the otherwise structured Swi5-Sfr1 ensemble (Fig. 2 and Supplementary Fig. 2). It was previously proposed that Swi5-Sfr1C inserts into grooves in the Rad51 nucleoprotein filament, locking the filament in an active confirmation^25, 26, 42^. While this could explain how Swi5-Sfr1 stabilizes the filament, it does not explain how Swi5-Sfr1 stimulates ATP hydrolysis and extensive strand transfer by Rad51, which are highly dynamic processes thought to involve dissociation of Rad51 from DNA^23^. It is possible that the flexibility of Sfr1N allows Swi5-Sfr1 to remain bound to Rad51 despite conformational changes in the filament. This would entail release of dissociating Rad51 molecules and re-binding of Sfr1N to molecules incorporated in the filament, thus preventing diffusion of Swi5-Sfr1 from the Rad51 filament. NMR interaction analysis suggested that binding of Sfr1N to Rad51 is relatively short-lived (see above), consistent with a model in which the rapid association and dissociation of Swi5-Sfr1 from Rad51 plays a role in stimulation of DNA strand exchange.

Although the role of Swi5-Sfr1 in promoting Rad51-dependent DNA repair is conserved in mammals^43, 45–49^, the precise mode of interaction with Rad51 appears to have undergone some divergence^43, 46, 50^. In *Saccharomyces cerevisiae*, the Swi5-Sfr1 homolog Sae3-Mei5 is produced only during meiosis and functions exclusively in the Dmc1 branch of meiotic HR^51, 52^. Both Swi5-Sfr1 and Sae3-Mei5 stimulate the strand exchange activity of Dmc1 via similar mechanisms^20, 22, 53, 54^, and although Sae3-Mei5 does interact directly with Rad51, it does not stimulate the activity of Rad51^55, 56^, unlike Swi5-Sfr1.

A consistent trend across all examined species is that Sfr1 plays some role in facilitating the interaction with the recombinase partner. Since the amino acid sequence of the N-terminal half of Sfr1 shows little conservation compared to the C-terminal half (Supplementary Fig. 4a), it is tempting to ascribe the similarities among species to the C-terminus. However, sequence divergence across large evolutionary distances does not necessarily reflect a lack of functional conservation for intrinsically disordered regions, which accumulate mutations at a higher rate than structured domains^57^. Notably, the large subunit of RPA contains an intrinsically disordered region whose function is conserved despite significant sequence divergence^58^, raising the possibility that the structure and/or function of Sfr1N is conserved. While empirical evidence is lacking, disorder predictions^59^ for the N-terminal half of *S. pombe* Sfr1 agree with the data presented here (Supplementary Fig. 7a) and similar profiles were generated for Sfr1 from *Schizosaccharomyces japonicus* and *Schizosaccharomyces octosporus* (Supplementary Fig. 7b,c). Furthermore, the N-terminal halves of mSfr1, hSfr1 and Mei5 are predicted to be enriched in intrinsically disordered regions (Supplementary Fig. 7d-f). This analysis also highlighted potential protein binding sites within the N-terminal halves of *S. japonicus* Sfr1, hSfr1 and Mei5. In agreement with this, the N-terminal half of Mei5 has already been shown to interact with Dmc1^52, 55^. Thus, in addition to the conserved function of Swi5-Sfr1 in promoting HR, the intrinsically disordered nature of Sfr1’s N-terminus and its role in facilitating interactions with recombinases may be evolutionarily conserved. Further studies will be required to test the validity of this prediction.

## ACKNOWLEDGEMENTS

We thank Tomohiro Koizumi for contributing to the early stages of this study; Yumiko Kurokawa and Yuki Ide for help with protein purification; and Ryoji Miyazaki, Hiroyuki Mori and Yoshinori Akiyama for help with the site-specific crosslinking experiments. This study was supported in part by Grants-in-Aid for Scientific Research on Innovative Areas (15H059749 to H.I., 18H04626 and 18H05426 to H. Takahashi), for Scientific Research (A) (18H03985 to H.I.), for Young Scientists (B) (17K15061 to B.A.), for Scientific Research (B) (18H02371 to H. Tsubouchi and 19H03160 to Y.M.), for Early-Career Scientists (19K16039 to K.I.), and a Research Fellowship (17J04051 to N.A.) from the Japan Society for the Promotion of Science. Y.M. also acknowledges support from the Takeda Science Foundation.

## AUTHOR CONTRIBUTIONS

B.A., M.S., H. Takahashi and H.I. conceived and designed the study. M.S. and M.K. performed all NMR experiments. B.A., N.A., K.I., and T.M. performed all other experiments. M.S., M.K. and H. Takahashi analyzed NMR data. B.A., N.A., K.I., T.M., S.K., Y.M., H. Tsubouchi, M.T. and H.I. analyzed all other data. S.K., Y.M., H. Tsubouchi, M.T., H. Takahashi and H.I. provided expertise and reagents. B.A., M.S., H. Takahashi and H.I. supervised the study. All authors contributed to writing the manuscript.

## DECLARATION OF INTERESTS

The authors declare no competing interests.

## ONLINE METHODS

### METHOD DETAILS

#### *S. pombe* and *E. coli* strains

The genotypes of *S. pombe* strains used in this study are listed in Table S1. Standard media was used for growth (YES), selection (YES with drugs or EMM), and sporulation (SPA), as described previously^60^. The genotypes of *E. coli* strains used in this study are listed in Table S2. Standard media was used for growth (LB) and selection (LB with antibiotics), unless otherwise indicated. All strains employed in this study are available upon reasonable request to the corresponding author (HI).

#### DNA damage sensitivity

A single colony was resuspended in 2 mL of YES and grown for 24 h (*rad^+^*) or 48h (*rad^−^*). Cells from these cultures were then seeded into 2 mL of fresh YES and grown for ∼14 h until they reached log phase (∼0.8 x10^7^ cells/mL). Cell density was adjusted to 2 x 10^7^ cells/mL, 10-fold serial dilutions were made, and 5 µL of each dilution was spotted onto YES plates (no treatment control) or YES plates containing the indicated genotoxins. In the case of UV irradiation, cells were spotted onto YES without any drug and treated with acute UV exposure of the indicated dose. The leftmost spot on each plate contains 1 x 10^5^ cells. Cells were photographed with a digital camera after the indicated growth period (2-4 days). For clonogenic assays, cells were grown as described above and spread onto several YES plates and irradiated with the indicated dose of UV. After 3 (*rad^+^*) or 4 (*rad^−^*) days of growth, colonies were counted.

#### Extraction of cellular proteins for immunoblotting

Cells (1 x 10^8^) were harvested and processed exactly as previously described^61^. Briefly, harvested cells were resuspended in 1mL of ice-cold water. 150µL of lysing solution (1.85 M NaOH 7.5% beta-mercaptoethanol) was added and mixed with the cells, followed by a 15 min incubation on ice. 150µL of 55% TCA was added, followed by a further 10 min incubation on ice. Precipitated proteins were pelleted by centrifugation (16,000 *g* 10 min 4°C) and dissolved with mixing (65°C 10 min) in 100 µL of urea loading buffer (8 M urea, 5% SDS, 200 mM Tris-Cl pH 6.8, 1 mM EDTA, 0.01% BPB, freshly supplemented with 10% volume each of 1 M DTT and 2 M Tris). Proteins were separated by SDS-PAGE, transferred to PVDF membranes, and detected with the indicated antibodies.

#### CD spectrometry

CD measurements were made using a Jasco J-720W spectrometer with a Peltier temperature controller. Sfr1N (4µM) in buffer N (20 mM sodium phosphate [pH 6], 25 mM NaCl, 1 mM DTT) was placed in a 0.1 cm path length quartz cuvette. The CD spectrum was acquired from 180 nm to 260 nm at 25 °C with a 1.0 nm bandwidth, 0.5 nm resolution, 50 nm/min scan speed, with 1 s averaging at each wavelength. Three spectra were averaged to give the spectrum of the protein and blank spectrum measured for buffer N alone was subtracted to produce the final spectrum.

#### NMR analysis of Sfr1N

Sfr1N (residues 1-176) was subcloned into pBKN220^20^, which was transformed into the *E. coli* strain BL21 (DE3) RIPL. Plasmids containing Sfr1N variants were prepared by using the protocol in the QuikChange Site-Directed Mutagenesis Kit (Agilent).

For the production of uniformly ^15^N-labeled or ^13^C and ^15^N-labeled proteins, *E. coli* cells were grown in M9 media supplemented with ^15^NH_4_Cl (1 g/L, Cambridge Isotope Laboratories) or with ^15^NH_4_Cl and ^13^C_6_-glucose (3 g/L, SHOKO Science), respectively. For the production of amino acid selectively labeled proteins, modified M9 media supplemented with a ^15^N-labeled amino acid (either Ala [200 mg/L]), Arg [200 mg/L], Ile [100 mg/L], Leu [100 mg/L], Lys [100 mg/L] or Phe [50 mg/L], all from Cambridge Isotope Laboratories) and other non-labeled amino acids (400 mg/L Ala, 400 mg/L Arg, 250 mg/L Asp, 50 mg/L Cys, 400 mg/L Glu, 400 mg/L Gly, 100 mg/L His, 100 mg/L Ile, 100 mg/L Leu, 150 mg/L Lys, 50 mg/L Met, 50 mg/L Phe, 150 mg/L Pro, 1000 mg/L Ser, 100 mg/L Thr, 50 mg/L Trp, 100 mg/L Tyr, and 100 mg/L Val) was used for culturing cells. Cultures were shaken in baffled flasks at 37 °C until the OD_600_ reached 0.8∼1.0. Protein expression was then induced by the addition of 0.5 mM isopropyl-β-D-thiogalactoside (IPTG) for 20 hours at 20 °C. For cultures with labeled amino acids, the induction was limited to ≤4 hours to minimize isotope scrambling. Sfr1N was then purified as described^24^, except the storage buffer used was buffer N (20 mM sodium phosphate [pH 6], 25 mM NaCl, 1 mM DTT). A truncated version of Sfr1N (127-176) was purified as a fusion protein with an N-terminal maltose-binding protein tag. Following purification with amylose resin (NEB), the tag was cleaved with Factor Xa and the cleaved peptide was isolated by ultrafiltration (Amicon Ultra-15 MWCO 10K, Merck) and subsequent solid phase extraction using a Sep-Pak C8 plus short cartridge (Waters).

NMR experiments were carried out using Bruker Avance III HD 500 and 800 spectrometers equipped with TCI cryoprobes at 25 °C. The spectra were processed using the program NMRPipe^62^ and analyzed using the program Sparky (Goddard, T. D. and Kneller, D. G. University of California, San Francisco).

For the main-chain resonance assignments of Sfr1N, ^13^C and ^15^N doubly labeled samples at concentrations of 0.15∼0.20 mM in buffer N mixed with 5% D_2_O were prepared and placed in 5 mm symmetrical microtubes (Shigemi). The ^1^H-^15^N HSQC spectrum^63, 64^ was acquired at a ^1^H frequency of 800 MHz with a scan number of 8, 1024 complex points and an acquisition time of 91.8 ms in the observed dimension, and 256 complex points and an acquisition time of 132 ms in the indirect dimension. The HNCA^65, 66^, HN(CO)CA^65, 66^, HNCACB^67^, HN(CO)CACB^68^, HNCO^65, 66^, and HN(CA)CO^69^ spectra were acquired at 800 MHz with a scan number of 8. All spectra were acquired with 512 complex points and an acquisition time of 45.9 m in the observed dimension. In the ^15^N dimension, all spectra except HNCACB were acquired with 45 complex points and an acquisition time of 23.1 ms, while the HNCACB spectrum was measured with 47 complex points and an acquisition time of 24.1 ms. The experiments with ^13^C_α_, ^13^C_α_/^13^C_β_, and ^13^CO evolution were acquired with 64, 128, and 43 complex points and acquisition times of 13.3, 10.6, and 8.90 ms in the ^13^C dimension, respectively.

To verify the signal assignments, the following samples were prepared and their ^1^H-^15^N HSQC spectra were obtained at 500 MHz; amino-acid selectively ^15^N-labeled (Ala, Arg, Ile, Leu, Lys, or Phe) Sfr1N(1-176), Arg-selectively ^15^N-labeled R97A variant of Sfr1N(1-176), Lys-selectively ^15^N-labeled K93A variant of Sfr1N(1-176), and uniformly ^15^N-labeled Sfr1 (127-176). From the 160 expected main-chain amide NH signals, 157 were detected (98%) and assigned to specific residues in Sfr1N. The remaining three signals from Q2, S3, and H51 could not be assigned due to line-broadening. NMR data were deposited in the Biological Magnetic Resonance Bank (BMRB) repository with accession number 27885.

#### Secondary structure analysis

The secondary structural elements were analyzed by calculating deviations of the observed ^13^C_α_, ^13^C_β_, and ^13^CO chemical shifts from their residue-dependent random coil values^30, 70^. Residues were deemed to form random coils if they displayed secondary chemical shift values within a limited range (between −0.7 and 0.7 for ^13^C_α_ and ^13^C_β_ atoms, and between −0.5 and 0.5 for ^13^CO atoms of non-proline residues, and −4 to 4 for all three ^13^C atoms of proline residues). The program TALOS+^31^ was also used to predict the secondary structural units where ^1^H_N_, ^13^C_α_, ^13^C_β_, ^13^CO, and ^15^N_H_ chemical shifts were used as input data. Predictions of disorder and protein binding sites for Sfr1 homologs were generated by DISOPRED3^59^.

#### Heteronuclear NOE

The heteronuclear {^1^H}-^15^N NOE experiment^32^ was carried out for uniformly ^15^N-labeled Sfr1N at 800 MHz. The NOE values were determined from the ratio I_NOE_/I_ref_, where I_NOE_ and I_ref_ indicate the signal intensities in the spectra acquired with and without 3 sec ^1^H presaturation, respectively. The NOE and reference spectra were acquired in an interleaved manner with a scan number of 32, 1024 complex points and an acquisition time of 91.8 ms in the observed dimension, and 256 complex points and an acquisition time of 132 ms in the indirect dimension.

#### NMR interaction analysis

250 μL of uniformly ^15^N-labeled Sfr1N at a concentration of 0.1 mM in buffer N was mixed with a 0.1 mM Rad51 solution in buffer N at Sfr1N:Rad51 molar ratios of 1:0, 1:0.25, 1:0.5, 1:0.75, and 1:1. These Sfr1N-Rad51 mixtures were concentrated to 250 μl (Amicon Ultra-4 MWCO 10K, Merck) and used for NMR measurements. For each mixture, ^1^H-^15^N HSQC spectra were acquired at 500 MHz with a scan number of 16, 1024 complex points and an acquisition time of 146 ms in the observed dimension, and 256 complex points and an acquisition time of 126 ms in the indirect dimension.

#### Site-specific crosslinking

*E. coli* strains used in this study are listed in Supplementary Table S1. Experiments were performed essentially as described^36^. Co-expression of Sfr1N (1-176, with or without a specific amber codon) and Rad51 was induced with 1 mM IPTG at an OD_600_ of ∼0.35 in *E. coli* strain BL21 (DE3) containing the pEVOL-pBpF plasmid either in the presence or absence of 1 mM *p*BPA. After 1 h at 30 °C, cultures were pre-chilled on ice for 5 min before 250 µL of cells were spotted in a radial manner onto petri dishes at 4 °C and UV irradiated for 10 min at a distance of 4 cm with a B-100 AP UV lamp (365 nm; UVP, LLC). 200 µL of cells was recovered from the plate and pelleted by centrifugation (20,000 *g* 5 min 4 °C). This pellet was dissolved in 70 µL of urea loading buffer^61^ and subjected to immunoblotting.

#### Purification of proteins for biochemical analyses

Previously published protocols were followed to purify Rad51^21^, RPA^20^, and Swi5-Sfr1 (wild type^20^, Sfr1N^24^, and Swi5-Sfr1C^24^). Swi5-Sfr1 mutants (3A, 4A, 7A) were purified by the same method as wild type Swi5-Sfr1 except they were diluted to ∼25 mM NaCl instead of ∼100 mM NaCl before being applied to the HiTrap Heparin column. All proteins were free of contaminating nuclease and ATPase activity for the duration of the relevant assays. In all assays where comparisons were made between reactions with and without protein, the equivalent volume of protein storage buffer was added instead of the protein.

#### Three-strand exchange assay

Strand exchange buffer (30 mM Tris-Cl [pH 7.5], 100 mM NaCl, 20mM KCl, 3.5 mM MgCl_2_, 2 mM ATP, 1 mM DTT, 5% glycerol, 8 mM phosphocreatine, 8 units/mL creatine phosphokinase) containing 10 µMnt PhiX174 virion DNA (NEB) was supplemented with 5 µM Rad51 and incubated at 37 °C for 10 min. The indicated concentration and variant of Swi5-Sfr1 was then added and the reaction was incubated for 10 min at 37 °C. Next, 1 µM of RPA was added and the reaction was incubated for 7 min at 37 °C. The reaction was initiated through the addition of 10 µMnt PhiX RF I DNA (NEB) linearized with ApaLI and incubated for a further 2 h at 37°C. The 10 µL reactions were subjected to psoralen-UV crosslinking to capture labile DNA structures and 1.95 µL of stop solution was added (30 mM Tris-Cl [pH 7.5], 44 mM EDTA, 3% SDS, 15 mg/mL proteinase K). Following a 15 min incubation at 37 °C, DNA was resolved in 1% agarose gels and stained with SYBR Gold (Thermo Fisher Scientific).

#### In vitro interaction assay

For the affi-gel interaction assay (Fig. 5B), BSA or Rad51 was covalently attached to Affi-gel 15 (2 µg protein/µL gel) according to the manufacturer’s instructions. 2 µg of the indicated Swi5-Sfr1 variant was diluted into 300 µL of Affi-gel buffer (25 mM HEPES-KOH [pH 7.5], 150mM NaCl, 3.5 mM MgCl_2_, 0.5 mM DTT, 0.05 µg/µL BSA, 10% glycerol, 0.05% igepal CA-630), input sample was taken, and 140 µL of this solution was mixed with 10 µL of Affi-BSA or Affi-Rad51. Reactions were then incubated with gentle mixing (30 °C 30 min). Following a brief centrifugation, flow-through samples were taken and the resin was washed with Affi-gel buffer (200 µL, x2). Bound proteins were eluted in 50 µL of SDS loading buffer with gentle mixing (37 °C 15 min). 9 µL (input, flow-through) or 3 µL (eluate) of sample was separated by SDS-PAGE, proteins were transferred to PVDF membranes and Sfr1 was detected with an anti-Sfr1 antibody^20^.

For the IP experiment in Supplemental Fig. S4C, 250 nM of a Swi5-Sfr1 variant and 250 nM of Rad51 were mixed on ice in 120 µL of IP buffer (30 mM Tris-Cl [pH 7.5], 150 mM NaCl, 3.5 mM MgCl_2_, 5% glycerol, 0.1% Igepal CA-630). Input sample was taken and 100 µL of the solution was incubated at 30 °C for 15 min. Protein A SureBeads (Bio-Rad) preincubated with anti-Rad51 antibody^20^ was added and mixtures were incubated with gentle mixing (4 °C 2 h). Beads were washed with IP buffer (500 µL, x3) and bound proteins were eluted in 50 µL of SDS loading buffer with gentle mixing (65 °C 10 min). Proteins were separated by SDS-PAGE, transferred to PVDF membranes and detected with the indicated antibodies.

#### Antibodies used for IP and immunoblotting

For the immunoblots of site-specific crosslinking experiments: anti-Rad51 (Rb 1:10,000; provided by Hiroshi Iwasaki); anti-Sfr1 (Rb 1:5,000; provided by Hiroshi Iwasaki). For the immunoblots of cellular proteins: anti-MYC (Rb 1:1,000; Sigma Aldrich C3956), anti-Rad51 (Rat 1:10,000; provided by Hiroshi Iwasaki), and anti-tubulin (Mu 1:10,000; Sigma Aldrich T5168). For the detection of Sfr1 in the affi-gel assay: anti-Sfr1 (Rb 1:5,000; provided by Hiroshi Iwasaki). For IP of Rad51 complexes, anti-Rad51 (Rb; provided by Hiroshi Iwasaki) was used, and for detection by immunoblotting: anti-Rad51 (Rat 1:10,000; provided by Hiroshi Iwasaki); anti-Sfr1 (Mu 1:1,000; provided by Hiroshi Iwasaki). Antibodies raised by our lab are available upon reasonable request to the corresponding author (HI).

#### Analysis of Rad51 filament dissociation kinetics

Anisotropy buffer (30 mM HEPES-KOH [pH 7.5], 100 mM KCl, 10 mM NaCl, 3 mM MgCl_2_, 1 mM ATP, 1 mM DTT, 5% glycerol) containing 1.5 µMnt of oligo dT (72 mer) with a 5’ TAMRA label was supplemented with 0.5 µM Rad51 and incubated at 25 °C for 5 min. Next, a Swi5-Sfr1 variant was added at the indicated concentration and the reaction was incubated at 25 °C for a further 5 min. This solution was transferred into a 0.3 x 0.3 cm quartz cuvette and the fluorescence anisotropy was monitored once per second for 60 seconds (25 °C, excitation 564 nm, emission 575 nm) to confirm filament formation. Next, a 1.0 x 1.0 cm quartz cuvette containing 2 mL of anisotropy buffer was placed into the spectrofluorometer with constant stirring (450 r.p.m) and, after 60 s of measurement, 50 µL of the solution containing Rad51 filaments with or without Swi5-Sfr1 was injected into this cuvette. Fluorescence anisotropy was then monitored once per second for the indicated time. Dissociation rate constants (*k_off_*) were calculated in KaleidaGraph. Cuvettes were purchased from Hellma Analytics.

#### ATPase assay

ATPase buffer (30 mM Tris-Cl [pH 7.5], 100 mM KCl, 20 mM NaCl, 3.5 mM MgCl_2_, 5% glycerol) containing 10 µMnt PhiX virion DNA was mixed on ice with 5 µM Rad51 and 0.25 µM of a Swi5-Sfr1 variant. Reactions were initiated through the addition of 0.5 mM ATP. Time zero was immediately withdrawn (10 µL) and mixed with 2 µL of 120 mM EDTA to terminate the reaction. Following incubation at 37 °C, aliquots were withdrawn at the indicated timepoints and processed as above. Upon completion of the time course, aliquots were diluted two-fold with water to reduce the concentration of ATP to 0.25 mM. Inorganic phosphate generated by ATP hydrolysis was then detected using a commercial malachite green phosphate detection kit (BioAssay Systems).

### QUANTIFICATION AND STATISTICAL ANALYSIS

#### DNA damage sensitivity assay

For the clonogenic survival assay, colonies were counted after 3 (*rad^+^*) or 4 (*rad^−^*) days of growth. The percentage of survival on the control plate (no UV treatment) was set to 100%. The expected number of colonies, based on the increased number of cells plated, was determined for the plates irradiated with the indicated dose of UV. The number of actual colonies was expressed as a fraction of the expected number, yielding the percentage of cells that survived the treatment. The values from three independent experiments were averaged and plotted, with the standard deviation of these averaged values depicted by error bars. In Fig. 7b, a one-way ANOVA followed by Tukey’s multiple comparisons test was performed. *, *P* < 0.05, n.s., not significant (*P* > 0.05). Further statistical information for Fig. 7b is listed below. Raw data is available upon reasonable request from the corresponding author (HI).

| Tukey's multiple comparisons test | Mean Diff. | 95.00% CI of diff. | Significant? | Summary | Adjusted P Value |
| --- | --- | --- | --- | --- | --- |
| *WT* vs. *sfr1d* | 23.8 | 13.4 to 34.21 | Yes | ** | 0.001 |
| *WT* vs. 7A | 10.16 | -0.2475 to 20.57 | No | ns | 0.0547 |
| *sfr1d* vs. 7A | -13.64 | -24.05 to -3.234 | Yes | * | 0.0163 |

| Test details | Mean 1 | Mean 2 | Mean Diff. | SE of diff. | n1 | n2 | q | DF |
| --- | --- | --- | --- | --- | --- | --- | --- | --- |
| *WT* vs. *sfr1d* | 25.14 | 1.337 | 23.8 | 3.392 | 3 | 3 | 9.92 | 6 |
| *WT* vs. 7A | 25.14 | 14.98 | 10.16 | 3.392 | 3 | 3 | 4.24 | 6 |
| *sfr1d* vs. 7A | 1.337 | 14.98 | -13.64 | 3.392 | 3 | 3 | 5.69 | 6 |

#### Three-strand exchange assay

Following staining with SYBR Gold (Thermo Fisher Scientific), gels were imaged using a LAS4000 mini (GE Healthcare). Densitometric analysis was performed using Multi Gauge software (version 3.2, Fujifilm) exactly as described^20, 21^. Background signal above the lds, NC and JM bands was subtracted from the corresponding values. The JM value was divided by 1.5 to compensate for the extra signal generated by these three-stranded DNA molecules. The sum of the values was set to 100%, and the percentage of total DNA corresponding to NC or JM was calculated. For the total yield, the percentage of DNA corresponding to NC and JM was combined. The values from three independent experiments were averaged and plotted, with the standard deviation of these averaged values depicted by error bars. Raw data is available upon reasonable request from the corresponding author (HI)

#### Analysis of Rad51 filament dissociation kinetics

To set a consistent end-point for the anisotropy graphs (Supplemental Fig. S4D-G), data were portrayed for the reactions until they reached a moving average (20 datapoints) of 0.106, which is equivalent to the value observed for DNA only. Dissociation rate constants (*k_off_*) were calculated in KaleidaGraph using the following equation: Anisotropy = (Amplitude of change in anisotropy) x e^(-koff x t)^ + (Minimum value of anisotropy) *k_off_* values from three independent experiments were averaged and plotted, with the standard deviation of these averaged values depicted by error bars. Raw data is available upon reasonable request from the corresponding author (HI)

#### ATPase assay

Absorbance values at 620 nm (A_620_) were obtained using a Nanodrop spectrophotometer (Thermo Fisher Scientific). A_620_ at time zero was subtracted from each value and these values were converted to concentrations of inorganic phosphate through the use of a standard curve. Graphs were plotted with inorganic phosphate concentration (µM) on the y-axis and time (min) on the x-axis. The gradient of the line of best fit was divided by the concentration of Rad51 to yield *k_cat_* values. *k_cat_* values from three independent experiments were averaged and plotted, with the standard deviation of these averaged values depicted by error bars. Raw data is available upon reasonable request from the corresponding author (HI).

## SUPPLEMENTARY INFORMATION

**Supplementary Table 1.**
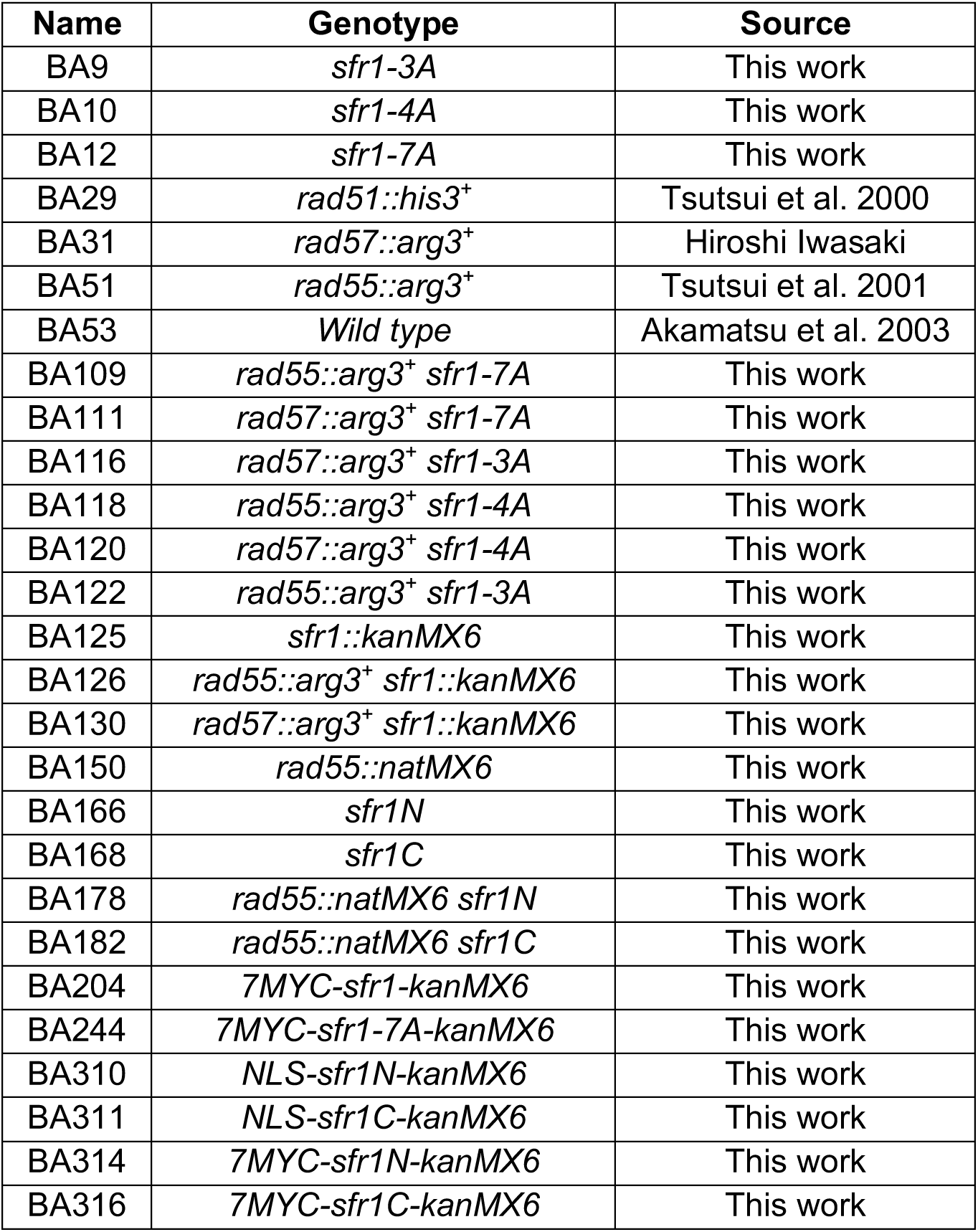
Genotypes of *S. pombe* strains are listed in the table below. All strains are isogenic derivatives of strain YA119 (Akamatsu et al. 2003) and are in the following genetic background: *Msmt-0 leu-1-32 ura4-D18 his3-D1 arg3-D1*

**Supplementary Table 2.** *E. coli* strains were constructed by transforming BL21 (DE3) containing the pEVOL-pBpF plasmid (Young et al. 2009) with a pBKN220 plasmid encoding Sfr1N (with or without a TAG mutation) and a pET28a plasmid encoding Rad51 (pBA33). Thus, strains only differ in the expression of Sfr1N or Rad51 and only these differences are indicated in the table below. A “–” sign indicates that cells were transformed with an empty vector whereas a “+” sign signals the presence of Sfr1N or Rad51 on that vector. If a codon in Sf1N was mutated to TAG, the mutated residue is listed instead of a “+” sign. Strains are listed in order of appearance.

| Name | Sfr1N / Rad51 |
| --- | --- |
| BA009 | – / + |
| BA0058 | – / – |
| BA0060 | + / – |
| BA0011 | + / + |
| BA0013 | K93 / + |
| BA0015 | K94 / + |
| BA0017 | R95 / + |
| BA0019 | A96 / + |
| BA0021 | R97 / + |
| BA0023 | E98 / + |
| BA0025 | A99 / + |
| BA0027 | K100 / + |
| BA0029 | N101 / + |
| BA0031 | I102 / + |
| BA0033 | L103 / + |
| BA0035 | L104 / + |
| BA0037 | K105 / + |
| BA0039 | P106 / + |
| BA0044 | F107 / + |
| BA0046 | K108 / + |
| BA0048 | S109 / + |
| BA0050 | P110 / + |
| BA0052 | L111 / + |
| BA0054 | R112 / + |
| BA0056 | Q113 / + |
| BA0062 | K157 / + |
| BA0064 | R158 / + |
| BA0066 | Q159 / + |
| BA0068 | K160 / + |
| BA0070 | R161 / + |
| BA0072 | L162 / + |
| BA0074 | F163 / + |
| BA0076 | K164 / + |
| BA0078 | S165 / + |
| BA0080 | P166 / + |

## SUPPLEMENTARY FIGURES

**Supplementary Figure 1.**
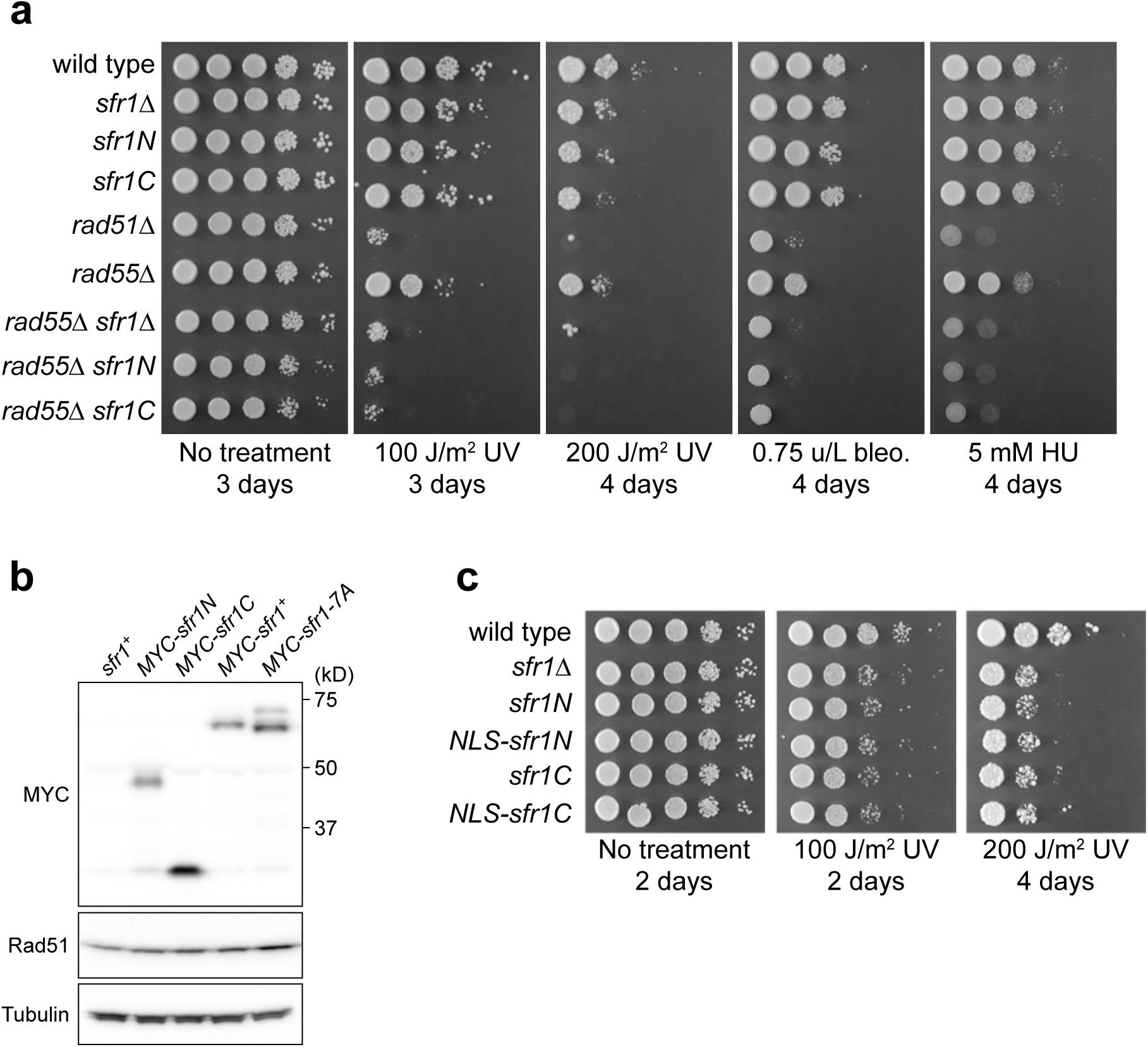
Sfr1N is essential for Swi5-Sfr1-dependent DNA repair. **(a,c)** DNA damage sensitivity of the indicated strains was assessed. NLS, nuclear localization signal of the SV40 large T antigen. Bleo., bleomycin. HU, hydroxyurea. Note that *sfr1Δ* does not sensitize cells to the indicated doses of bleo. or HU unless Rad55-Rad57 is absent. **(b)** Proteins were extracted from log phase cultures, separated by SDS-PAGE and probed with the indicated antibodies. Tubulin serves as a loading control.

**Supplementary Figure 2.**
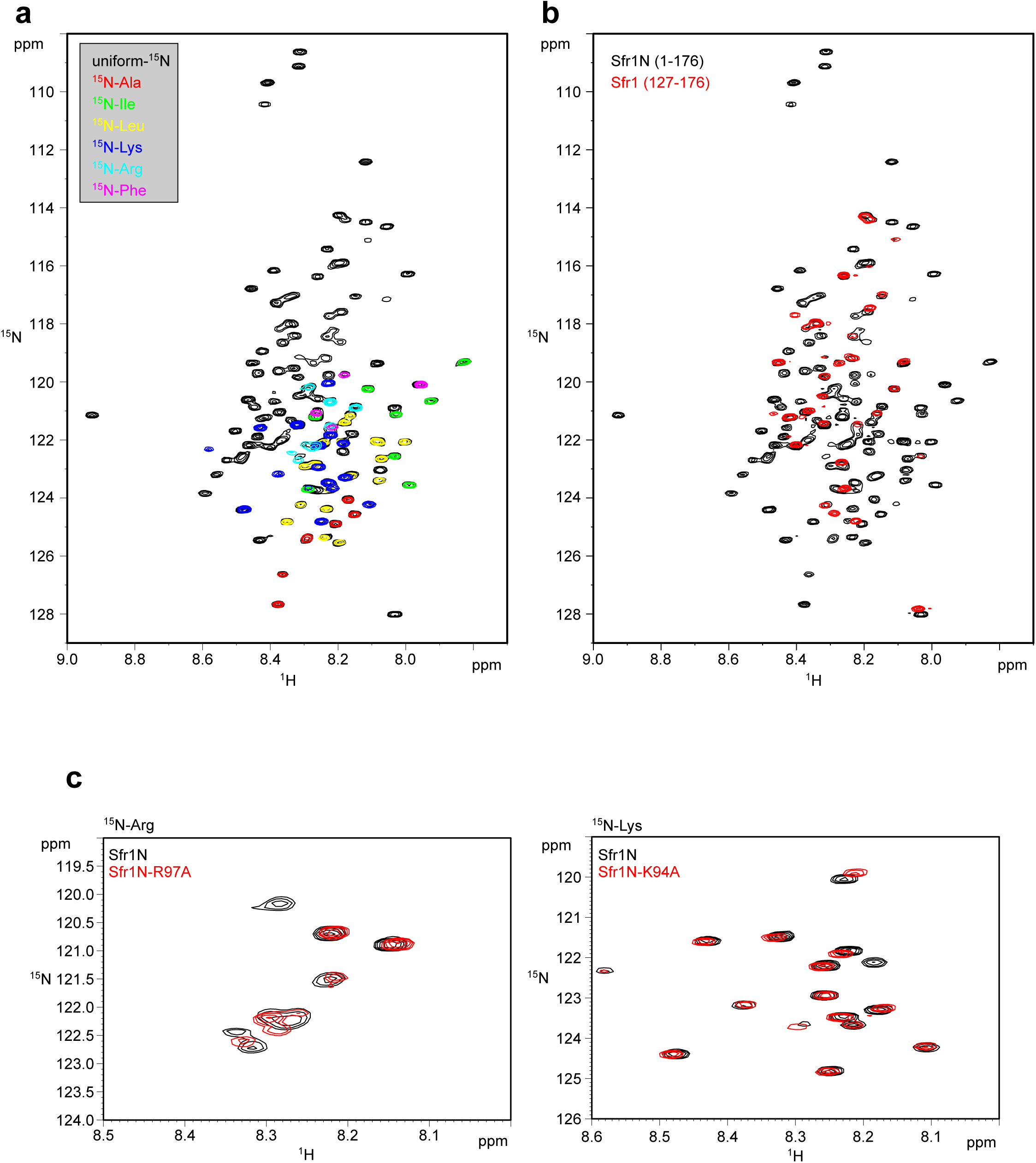
Validation of Sfr1N signals by ^15^N labelling. **(a)** Superimposed ^1^H-^15^N HSQC spectra of Sfr1N with the indicated ^15^N-labelling. **(b)** Superimposed ^1^H-^15^N HSQC spectra of uniformly ^15^N-labeled Sfr1N and a truncated variant, Sfr1 (127-176). **(c)** Left, superimposed ^1^H-^15^N HSQC spectra of Arg-selectively ^15^N-labeled Sfr1N or Sfr1N-R97A. Right, superimposed ^1^H-^15^N HSQC spectra of Lys-selectively ^15^N-labeled Sfr1N or Sfr1N-K94A.

**Supplementary Figure 3.**
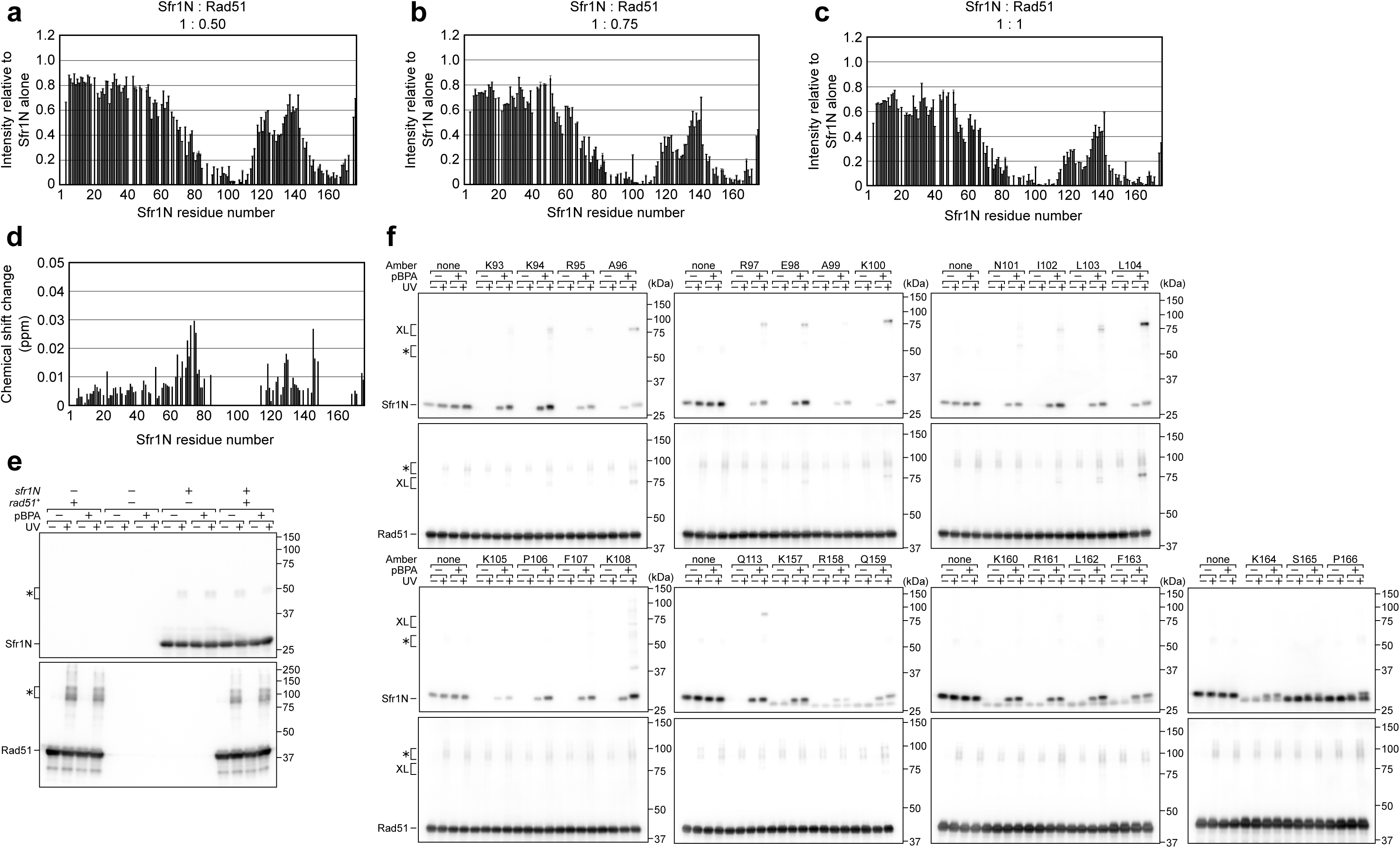
Changes in Sfr1N NMR signal intensity in response to Rad51 titration and site-specific crosslinking of Sfr1N to Rad51. **(a-c)** NMR signal intensity of Sfr1N in the presence of the indicated ratio of Rad51 relative to the intensity observed in the absence of Rad51. **(d)** Chemical shift changes observed for the main chain amide NHs of Sfr1N induced by the addition of equimolar Rad51. Chemical shift changes were calculated using the formula Δδ = [(Δδ*_H_*)^2^ + (Δδ*_N_* × 0.18)^2^]^1/2^, where Δδ_H_ and Δδ_N_ represent the ^1^H and ^15^N chemical shift differences, respectively. The residues that were severely attenuated by the addition of Rad51 were not analyzed and are shown by blanks. **(e)** The extent of non-specific crosslinking was assessed by individual and co-expression of Sfr1N and Rad51 in *E. coli*. **(f)** Specific crosslinking was assessed by expressing Rad51 with a version of Sfr1N in which a residue of interested was mutated to TAG, which encodes the photoreactive amino acid *p*BPA in this system.

**Supplementary Figure 4.**
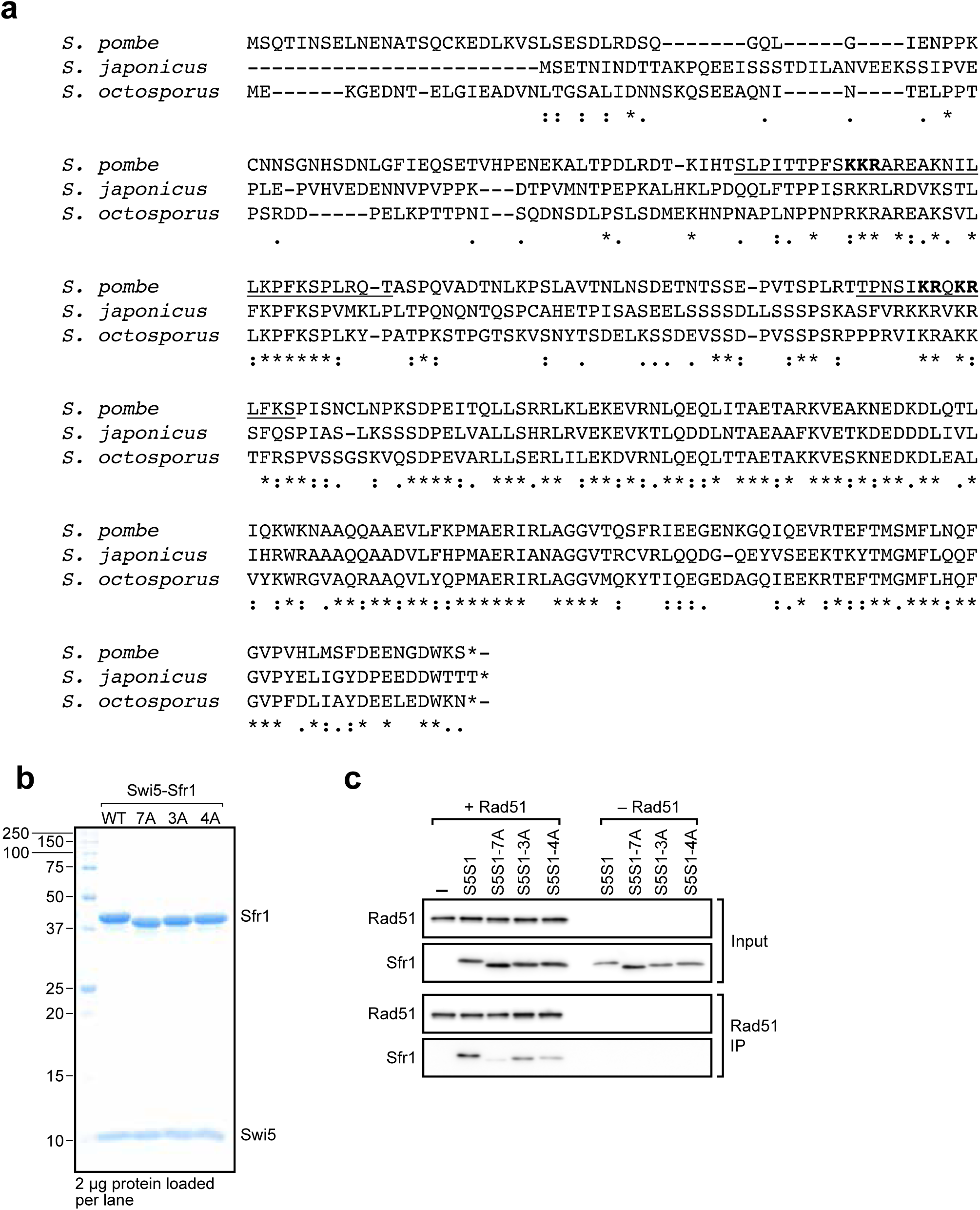
Identification of residues within Sites 1 and 2 that are important for the functional interaction with Rad51. **(a)** Sequence alignment of Sfr1 homologs from the genus *Schizosaccharomyces* were generated with Clustal Omega. Sites 1 and 2 in *S. pombe* Sfr1 are underlined. Residues shown in bold are consecutive, positively charged residues within Sites 1 and 2 that were targeted for mutation. **(b)** CBB-stained gel of purified Swi5-Sfr1 (wild type mutants). 7A has mutations in all bold residues from **(a)**, whereas 3A and 4A only contain mutations in bold residues from Sites 1 and 2, respectively. Size markers are annotated on the left in kD. Note that Sfr1 migrates slower than expected from its calculated molecular weight (∼33.6 kD), as previously reported^20^. This property is attributable to its N-terminal half^25^, hence why mutation of residues in Sites 1 and 2 affect its electrophoretic mobility. **(c)** Swi5-Sfr1 (S5S1, wild type or mutants) was mixed with Rad51, complexes were IP’d with anti-Rad51 antibodies and the contents of these IPs were assessed by immunoblotting. “–” indicates the omission of Swi5-Sfr1.

**Supplementary Figure 5.**
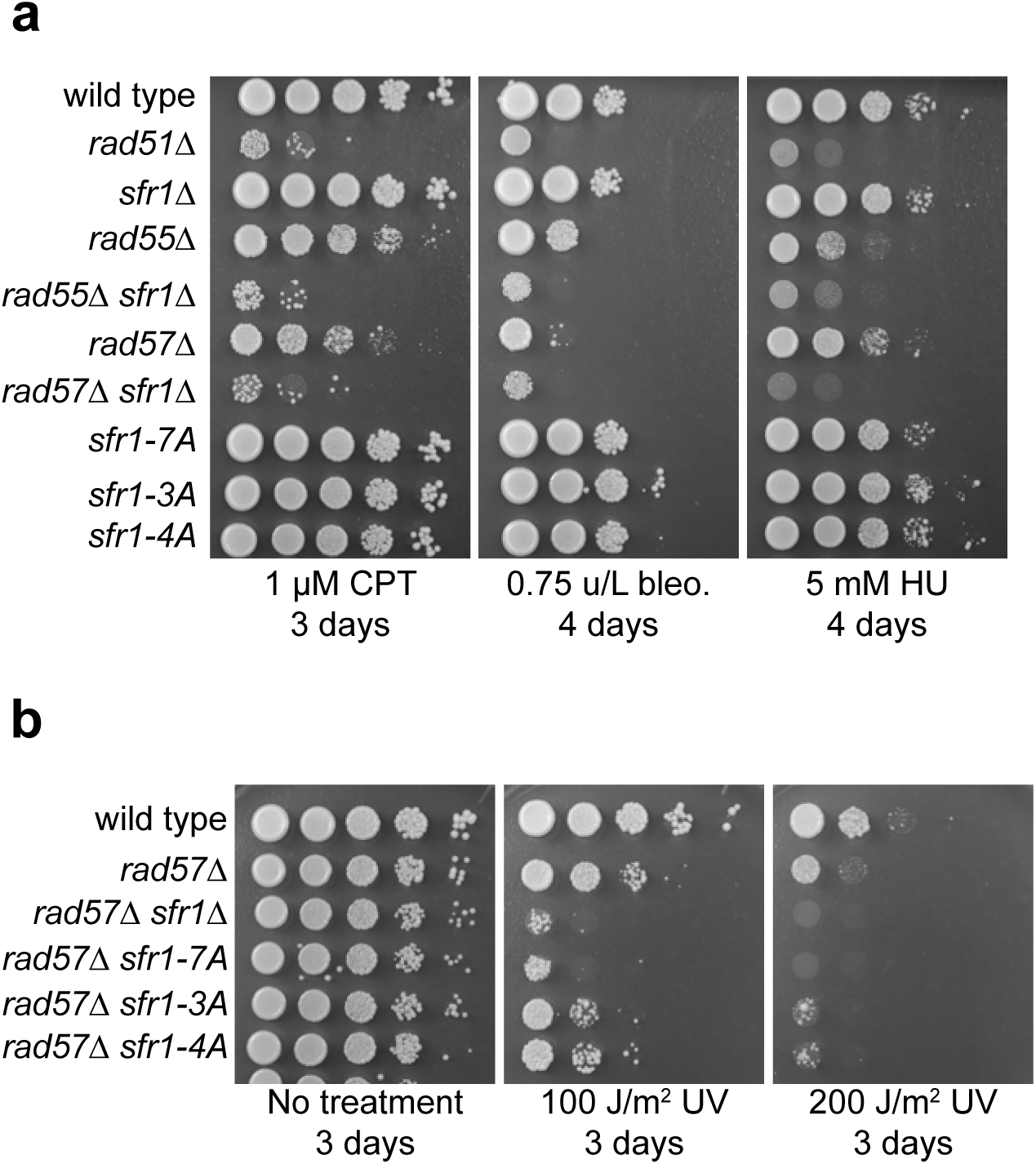
DNA repair in Rad51 interaction mutants is dependent on Rad57. **(a,b)** DNA damage sensitivity of the indicated strains was assessed. “No treatment” plate for **(a)** is shown in Fig. 7a. CPT, camptothecin. Bleo., bleomycin. HU, hydroxyurea. Note that *sfr1Δ* does not sensitize cells to the indicated doses of CPT, bleo. or HU unless Rad55-Rad57 is absent.

**Supplementary Figure 6.**
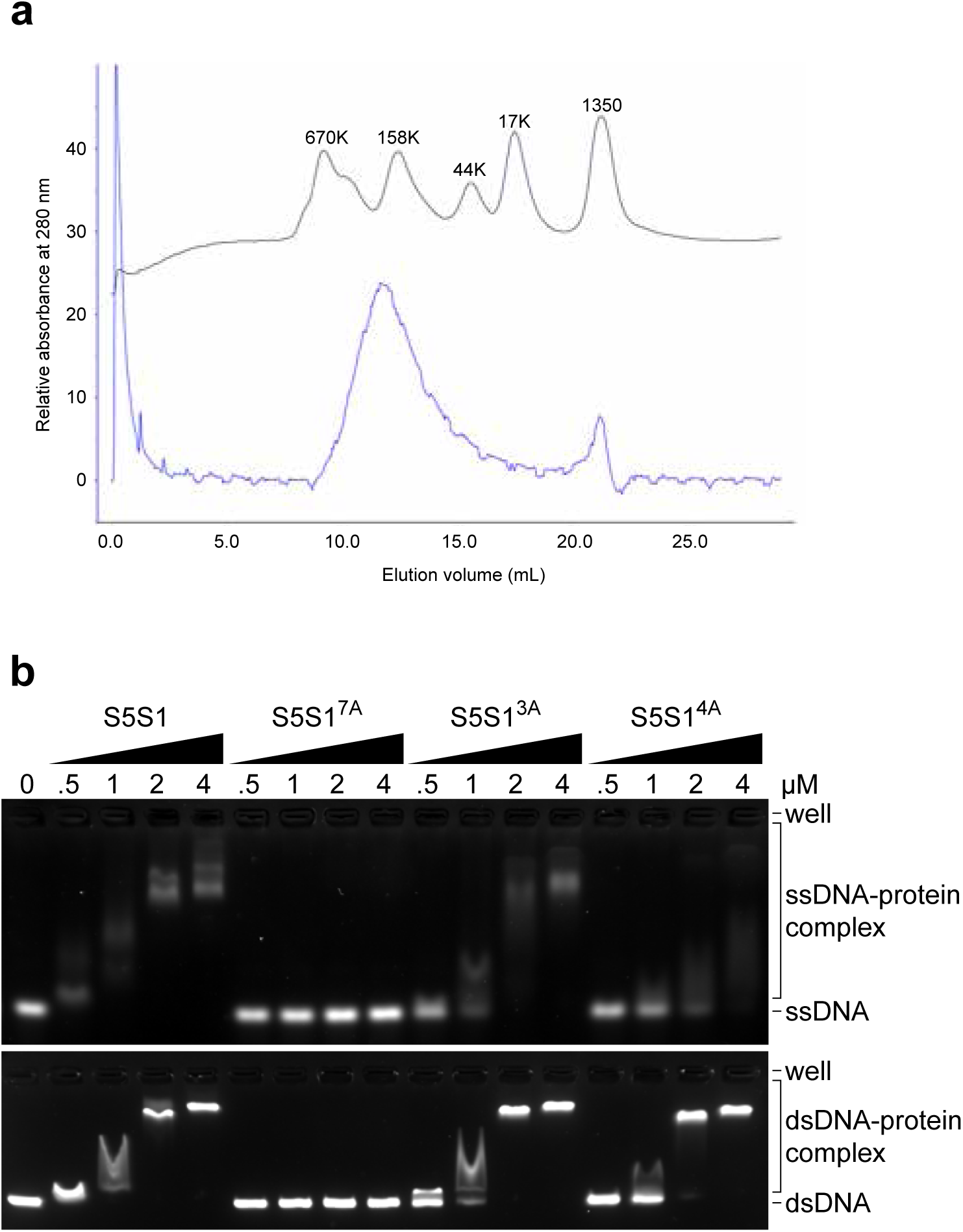
Molecular sizing of Rad51 and DNA binding by Swi5-Sfr1. **(a)** Rad51 was analyzed by size-exclusion chromatography. The blue and navy lines indicate the elution profiles of Rad51 and molecular weight standards, respectively. **(b)** Swi5-Sfr1 (S5S1, wild type or mutants) was incubated with ssDNA (top) or dsDNA (bottom) and protein-DNA complexes were resolved by agarose gel electrophoresis.

**Supplementary Figure 7.**
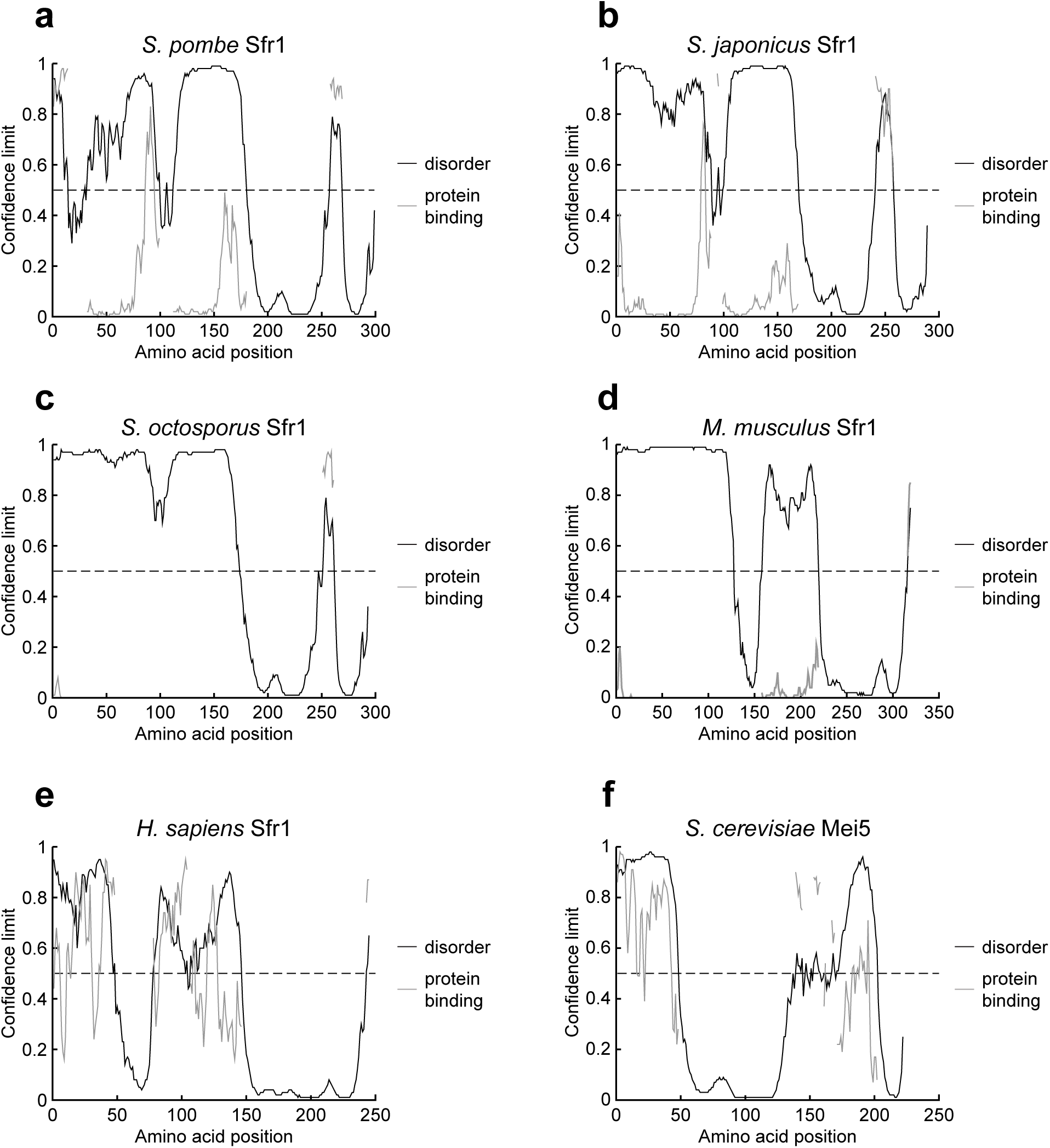
N-terminal intrinsically disordered domains may be an evolutionarily conserved feature of Sfr1. **(a-f)** DISOPRED3 was used to predict the likelihood of disorder and protein binding in orthologs of Sfr1 from the indicated species. The likelihood of disorder is shown on the y-axis and residues of Sfr1 are displayed on the x-axis. The dashed line represents a confidence limit of 0.5, above which point DISOPRED3 predicts that a residue is more likely to be disordered than structured. Sequences were obtained from UniProt (*S. pombe*, Q9USV1; *S. japonicus*, B6JYP3; *S. octosporus*, S9R1E2; *Mus musculus*, Q8BP27; *Homo sapiens*, Q86XK3; *S. cerevisiae*, P32489).

